# A Physcomitrella PIN protein acts in spermatogenesis and sporophyte retention

**DOI:** 10.1101/2022.07.05.498815

**Authors:** Volker M. Lüth, Christine Rempfer, Nico van Gessel, Oliver Herzog, Melanie Hanser, Marion Braun, Eva L. Decker, Ralf Reski

**Affiliations:** Plant Biotechnology, Faculty of Biology, University of Freiburg, Freiburg, Germany; Spemann Graduate School of Biology and Medicine (SGBM), University of Freiburg, Freiburg, Germany; CIBSS – Centre for Integrative Biological Signalling Studies, University of Freiburg, Freiburg, Germany; Cluster of Excellence livMatS @ FIT – Freiburg Center for Interactive Materials and Bioinspired Technologies, University of Freiburg, Freiburg, Germany

**Keywords:** auxin, bryophyte, flagellum, moss, plant development, sperm, spermatozoid

## Abstract

- The auxin efflux PIN-FORMED (PIN) proteins are conserved in all land plants and important players in plant development. In the moss Physcomitrella (*Physcomitrium patens*) three canonical PINs (PpPINA-C) are expressed in the leafy shoot (gametophore). PpPINA and PpPINB show functional activity in vegetative growth and sporophyte development. Here, we examined the role of PpPINC in the life cycle of Physcomitrella.
- We established reporter and knockout lines for PpPINC and analysed vegetative and reproductive tissues using microscopy and transcriptomic sequencing of moss gametangia.
- PpPINC is expressed in immature leaves, mature gametangia and during sporophyte development. The sperm cells (spermatozoids) of *pin*C knockout mutants exhibit increased motility and an altered flagella phenotype. Further, the *pin*C mutants have a higher portion of differentially expressed genes (DEGs) related to spermatogenesis, increased fertility, and an increased abortion rate of premeiotic sporophytes.
- Here, we show that PpPINC is important for spermatogenesis and sporophyte retention. We propose an evolutionary conserved way of polar growth during early moss embryo development and sporophyte attachment to the gametophore, while suggesting the mechanical function in sporophyte retention of a ring structure, the Lorch ring.

## Introduction

The auxin signal transduction pathway is conserved in all land plants (Paponov *et al*., 2009; Flores-Sandoval *et al*., 2015; Thelander *et al*., 2018; Cancé *et al*., 2022). In the model moss Physcomitrella (*Physcomitrium patens*), the main auxin biosynthesis pathway is, as in Arabidopsis, the conversion of tryptophan by TAR enzymes to indole-3-pyruvate (IPyA) from where it is converted by YUC enzymes into active auxin (indole-3-acetic acid, IAA) (Landberg *et al*., 2020). Auxin homeostasis plays an important role during the life cycle of Physcomitrella, maintaining growth and organogenesis (Ludwig-Müller *et al*., 2009; Thelander *et al*., 2018). However, there is no indication for a polar auxin transport in the leafy moss shoot, the gametophore (Fujita *et al*., 2008), although the protein family responsible for polar auxin transport, PIN-FORMED (PIN), is conserved in all land plants, including bryophytes (Bennett *et al*., 2014a; Zhang *et al*., 2020a). Instead, auxin transport appears to be diffusive in the moss gametophore (Coudert *et al*., 2015).

Canonical PIN proteins share four highly conserved motifs in the hydrophilic loop and a strong conservation in the N’- and C’-terminal transmembrane regions, while noncanonical PINs are defined by a higher variability (Bennett *et al*., 2014a). Structures and mechanism of auxin transport have recently been elucidated in great detail for the *Arabidopsis thaliana* PIN8 protein (Ung *et al*., 2022). The Physcomitrella genome encodes four PIN genes, the three canonical *PpPINA*, *PpPINB* and *PpPINC*, and the noncanonical *PpPIND*, with *PpPINA* and *PpPINB* being the most similar to each other (Bennett *et al*., 2014b). The Physcomitrella PIN proteins form an outgroup with other bryophytes to vascular plants, where the canonical PpPINs cluster with PIN proteins from other mosses, while the noncanonical PpPIND is separated together with several PIN proteins of the liverwort *Marchantia polymorpha* (Bennett *et al*., 2014a).

In the gametophore, auxin accumulates beneath the stem apex and at the stem base (Bierfreund *et al*., 2003; Fujita *et al*., 2008). Apical stem cells show the highest expression of *PpTAR* genes, indicating biosynthesis of IPyA (Landberg *et al*., 2020). The growth of developing leaves (phyllids) in Physcomitrella is marked by high auxin-signalling related activity (Thelander *et al*., 2019) with all three canonical *PpPIN* genes being active (Viaene *et al*., 2014). While *PpPINA* is the highest expressed *PIN* gene in Physcomitrella tissues, *PpPINC* has the lowest expression level of all canonical PINs (Bennett *et al*., 2014b). Single knockouts of *PpPIN* genes have no severe effect on gametophore growth (Viaene *et al*., 2014), while the double knockout of *PpPINA* and *PpPINB* leads to elongated phyllids, a phenotype similar to Physcomitrella gametophores treated with excess auxin or auxin transport inhibitors (Decker *et al*., 2006; Bennett *et al*., 2014b).

When introduced to short day conditions with low temperatures, the monoecious moss Physcomitrella initiates the formation of sexual organs on the gametophore apex (Hohe *et al*., 2002) with specialized stem cells for female archegonia and male antheridia development (Kofuji & Hasebe, 2014). Growth of antheridia and archegonia is highly synchronized, beginning with the formation of the antheridia while archegonia develop later but mature faster, so that self-fertilization is possible (Cove, 2005; Landberg *et al*., 2013). The development and growth of gametangia is controlled by the auxin-biosynthesis regulators *SHORT INTERNODE/STYLISH* (*SHI/STY*) and TAR enzymes, influencing the neck length of archegonia and growth of antheridia. During antheridia development, the expression of *PpSHI* and *PpTAR* genes overlap with the expression of *PpPINA* and the accumulation of auxin in apical cells, before spermatogenesis begins. This activity slowly declines and reaches its lowest point during spermatogenesis, indicating a process where auxin plays a minor role (Landberg *et al*., 2013; Landberg *et al*., 2020). Spermatogenesis in Physcomitrella is tightly regulated, producing motile, biflagellate sperm cells (spermatozoids), relying on the availability of water to swim to the egg cell (Reski, 1998; Cove, 2005; Ortiz-Ramírez *et al*., 2017; Koshimizu *et al*., 2018; Gu *et al*., 2022). As in antheridia, the activity of auxin biosynthesis, signalling and accumulation are the highest in archegonia during early development. The precursor egg cell and apical neck cells show the highest activity, while there is minimal expression during egg maturation (Landberg *et al*., 2013; Landberg *et al*., 2020).

After fertilization of the egg cell, the zygote develops into a diploid embryo, resulting in the sporophyte (Horst *et al*., 2016). The Physcomitrella sporophyte is reduced compared to other mosses (Kirbis *et al*., 2020), even within its own family (Ostendorf *et al*., 2021), and consists of the sporophyte foot, a short seta and the spore capsule, which rips open after maturation to release the haploid spores (Cove, 2005). Auxin is distributed dynamically in a polar manner during sporophyte development, with an auxin maximum in the apex of the early embryo which later localizes to the foot of the young sporophyte where it slowly recedes during maturation (Fujita *et al*., 2008). The sporophyte foot is secured in a maternal cavity (vaginula) and covered at its base with haustorial cells, important for nutrient uptake from the gametophore (Regmi *et al*., 2017). Sporophyte formation is complex and regulated by a number of genetic elements (Sakakibara *et al*., 2008; Mosquna *et al*., 2009; Horst *et al*., 2016; Ortiz-Ramírez *et al*., 2016; Hashida *et al*., 2020; Kirbis *et al*., 2020; Takechi *et al*., 2021, Landberg *et al*. 2022; Lopez-Obando *et al*., 2022). However, auxin remains a crucial player in sporophyte development (Fujita *et al*., 2008) through the function of the two Physcomitrella PpPINA and PpPINB proteins (Bennett *et al*., 2014b), whereas the role of the third canonical PIN, PpPINC, remains unclear.

Here, we elucidate the role of PpPINC in the Physcomitrella life cycle. We found that PpPINC influences spermatogenesis-related gene expression, controls motility and morphology of moss sperm cells, and is important for the retention of the diploid sporophyte on the haploid gametophyte, and thus preventing early abortion of premeiotic sporophytes, while it has no obvious role in the development of the gametophyte, the protonemata and gametophores.

## Material and Methods

### Plant material and culture conditions

The *Physcomitrella patens* (Hedw.) Bruch & Schimp. ecotype Gransden covers several laboratory strains which are descendants of the first original cultivated single clone (Haas *et al*., 2020) and was recently renamed to *Physcomitrium patens* (Hedw.) Mitt.. We used as wild type (WT) a fertile Gransden line, which underwent sexual reproduction regularly as a basis for all transgenic lines. Plants were cultivated using Knop medium (pH 5.8) according to Reski & Abel (1985) containing microelements according to Egener *et al*. (2002). For solid medium, 12 g/l agar (OXOID, Thermo Scientific) were added. Standard growth conditions were long day 16 hours light with 70±5 µmol m^-2^ s^-2^ at 22°C. Sporophyte induction was modified after Hohe *et al*. (2002). Plants were grown in long day conditions, before being transferred to sporophyte inducing conditions (Hohe *et al*., 2002). At day 18 of sporophyte induction, plants were watered (H_2_O dest., 10 ml for 9 cm petri dish), which was removed from the plate at day 25. All moss lines used are stored in the International Moss Stock Center (IMSC; https://www.moss-stock-center.org) with the following accession numbers: *PinCPromCit =* 40917, *pin*C#10 = 40918, *pin*C#29 = 40919, *pin*C#69 = 40420, WT = 40095.

### Generation of transgenic lines

Transgenic lines were created via highly efficient homologous recombination (Hohe *et al*., 2004) in transformed protoplasts (Hohe & Reski, 2002). For the generation of targeted *pin*C mutants the region upstream from the beginning of the first exon to the untranslated region after exon six was amplified (2915 bp) from genomic DNA using the following primers: P3-KO Fw + P3-KO Rv (Supplemental Table S1). The amplified fragment was sub-cloned into the vector *pJET1.2*. Using *SacI* and *NcoI* a fragment of the *PpPINC* gene (1696 bp) was replaced with a sulfadiazine selection cassette (Parsons *et al*., 2012). For the generation of the transcriptional reporter line, 2.1 kb upstream of the start codon of the *PpPINC* gene were fused to the citrine CDS and nos terminator via Gibson cloning (Gibson *et al*., 2009), using primers 5PinCprom_f+CA5 + 3PinCprom_r+Citrin (Supplemental Table S1). The expression cassette was cloned between homologous regions of the carbonic anhydrase locus (Wiedemann *et al*., 2018), erasing the citrine expression of the parental plant when correctly integrated into the genome.

### Molecular analysis of transgenic lines

Initial screening of knockout and reporter lines was done with leaflet PCR according to Schween *et al*. (2002), to test for the presence of the construct. For RT-PCR total RNA was extracted from 6-weeks-old gametophores, 21 days after the sporophyte induction started using the innuPREP Plant RNA Kit (Analytik Jena AG, Jena, Germany) and reversely transcribed using oligo-d(T)16 primers with Superscript III reverse transcriptase (Life Technologies, Thermo Fisher Scientific). Analysis for the absence of *PpPINC* RNA was performed with the gene-specific primers Pin3f_ex1-2 + Pin3r_ex2 (Supplemental Table S1). Presence of cDNA was tested with C45_fwd and C45_rev (Supplemental Table S1) amplifying the constitutively expressed gene L21. Transgene copy numbers were tested via quantitative Real-Time PCR according to Noy-Malka *et al*. (2014). Genomic DNA was isolated from protonema one week after the last tissue disruption using the innuPREP Plant DNA Kit (Analytic Jena AG). Transgene copy numbers were determined comparing relative values of the transgene 35S promoter (35SPqPCR_f + 35SPqPCR_r; Supplemental Table S1) with the single copy transgene carbonic anhydrase line used in Wiedemann *et al*. (2018), for normalization the single copy gene CLF was used (Noy-Malka *et al*., 2014).

### Tissue isolation for gene expression analysis using SMARTseq

Triplicates of tissue samples were collected 18 days after gametophores were exposed to sporophyte-inducing conditions. Mature archegonia and antheridia were collected manually using a stereoscope (Olympus SZX7) and stored directly in TRIzol® (Fisher Scientific GmbH, Schwerte, Germany). All lines used were grown together on the same plate. For each sample eight archegonia or antheridia were collected. Tissues were homogenized using small pistils and mixed with chloroform. The aqueous phase was then processed using the Direct-zol RNA Microprep Kit (Zymo Research Europe GmbH, Freiburg, Germany). The resulting RNA was treated with RiboLock RNase Inhibitor (Fisher Scientific GmbH, Schwerte, Germany). The cDNA library was created at the Genomics Unit in the Instituto Gulbenkian de Ciencia, Portugal according to Picelli *et al*. (2014). Libraries were sequenced using the Illumina RNAseq platform from Novogene (Novogene Company Limited, Cambridge, UK).

### Transcriptomic data processing

Raw data was trimmed using Trim Galore (Version 0.6.6; adapter stringency = 1 bp; minimum required sequence length for retaining a read pair = 20 bp; 3’clipping = 1 bp). All further steps were done using the Galaxy platform (Afgan *et al*., 2018). The 150 bp paired-end reads were then mapped to the Physcomitrella genome version 3.3 downloaded from Phytozome (Goodstein *et al*., 2012) using HISAT2 (Galaxy Version 2.1.0; spliced alignment activated). Mapped reads were counted with feature counts (Galaxy Version 2.0.1; excluding chimeric fragments; only fragments with both reads aligned, GFF feature type filter = CDS; GFF gene identifier = gene_id). Differential gene expression was analyzed using DESeq2 (Galaxy Version 2.11.40.6+galaxy1) and filtered for enriched gene ontology (GO) terms with GOEnrichment (Galaxy Version 2.0.1). Quality of mapping and read counts were controlled with MultiQC (Galaxy Version 1.11+galaxy0) and FastQC (Galaxy Version 0.73+galaxy0).

### Calcium measurement in sperm cells

Sperm packets of single antheridia where extracted directly after being released from the antheridium and placed on polylysine-covered glass slides according to Horst & Reski (2017). Sperm cells where incubated in a Fluo-4 solution according to Ortiz-Ramírez *et al*. (2017) for 20 minutes, photographed (see microscopy) and pictures were then analysed using ImageJ (Schneider *et al*., 2012).

### Spermatozoid motility assessment

Mature antheridia were prepared on the moss apex, placed under a glass slide and sealed with nail polish to prevent water evaporation. Ripe antheridia opened without applying pressure, releasing sperm packets, which were then analyzed for motility.

### Sporophyte count

For all lines we analysed all gametophores on the respective plate with three plates for each line. Sporophytes were counted 4 weeks after watering (see sporophyte induction culture conditions). To account for sporophytes, embryos, empty vaginulae and aborted sporophytes we removed the phyllids of the gametophore apices.

### Statistical analysis

Stem length, leaf length and leaf width were tested with Student’s t-Test with p < 0.05. Significance of motility was assessed with one-sided ANOVA and Tukey-Kramer test, flagellar phenotype with one-sided ANOVA. All tests were performed with Microsoft Excel and the XLMiner Analysis ToolPak.

### Protein alignments and motif analysis

Needleman-Wunsch (Needleman & Wunsch, 1970) and multiple sequence alignments (Larkin *et al*., 2007) with Clustal Omega (1.2.4) were performed using the European Bioinformatics Institute (EMBL-EBI) web tools (https://www.ebi.ac.uk/services) (Madeira *et al*., 2019). Needleman-Wunsch alignments were performed with BLOSUM62, a gap penalty of 10 and extend penalty of 0.5. Multiple sequence alignments were performed with default settings.

### Flow cytometry

Flow cytometry analysis was performed according to Heck *et al*. (2021) to determine ploidy of the transgenic lines (*pin*C#10, *pin*C#29, *pin*C#69, *pinCPromCit*). As there is a risk of creating diploid plants during PEG-mediated transformation of protoplasts, which could alter gene expression compared to haploid plants (Rempfer *et al.,* 2022), only haploid transgenics were analysed further.

### Microscopy

For preparation of samples, we used an Olympus SZX7 stereoscope and for extraction of sperm packages a Zeiss Axiovert microscope. Fluorescence and bright-field microscopy pictures were taken with a Zeiss Axioplan 100 with a Zeiss MRc5 camera and Zeiss AxioVision software (Version 3.8.2).

### Phylogenetic reconstruction and transmembrane prediction

Based on reference sequences from *A. thaliana* and *P. patens* we conducted proteome-wide sequence searches with DIAMOND (v0.9.19, Buchfink *et al*., 2021) against *Anthoceros angustus* (Zhang *et al*., 2020b), *Calohypnum plumiforme* (Mao *et al*., 2020), *Chlamydomonas reinhardtii* (Merchant *et al*., 2007), *Ceratodon purpureus* (Carey *et al*., 2021), *Ceratopteris richardii* (Marchant *et al*., 2022), *Funaria hygrometrica* (Kirbis *et al*., 2021), *Klebsormidium nitens* (Hori *et al*., 2014), *Marchantia polymorpha* (Bowman *et al*., 2017), *Mesotaenium endlicherianum* (Cheng *et al*., 2019)*, Oryza sativa* (Ouyang *et al*., 2006), *Picea abies* (Nystedt *et al*., 2013), *Populus trichocarpa* (Tuskan *et al*., 2006), *Selaginella moellendorffii* (Banks *et al*., 2011), *Sphagnum fallax* (v1.1, DOE-JGI, http://phytozome.jgi.doe.gov/*), Sphagnum magellanicum* (v1.1, DOE-JGI, http://phytozome.jgi.doe.gov/*), Spirogloea muscicola* (Cheng *et al*., 2019), and *Vitis vinifera* (Jaillon *et al*., 2007). Multiple sequence alignment of protein sequences was performed with UPP/SEPP (v4.5.1, Nguyen *et al*., 2015) and converted into a codon-aware alignment of coding sequences with PAL2NAL (v14, Suyama *et al*., 2006). A maximum-likelihood tree was calculated with RAxML (v8.2.12, Stamatakis, 2014) using the “GTRGAMMA” model and 1,000 bootstrap replicates and visualized using R (v4.2.1, R Core Team 2022) and the ggtree package (v3.4.4, Yu *et al*., 2017) alongside transmembrane domains which were predicted by DeepTMHMM (v1.0.15, Hallgren *et al*., 2022).

## Results

### Physcomitrella PIN family

The Physcomitrella genome (Lang *et al*., 2018) encodes three canonical PIN proteins, PpPINA (Pp3c23_10200), PpPINB (Pp3c24_2970) and PpPINC (Pp3c10_24880). These proteins are similar in structure and length (Figure 1), consisting like all canonical land plant PINs of two transmembrane regions (five helices each), separated by a hydrophilic loop (Figure 1). The transmembrane regions have the same length in all three proteins, whereas the hydrophilic loop of PpPINC is 15 amino acids (AA) shorter than those of PpPINA and PpPINB (Table 1). While PpPINA and PpPINB are very similar in their AA sequences, PpPINC differs more, especially in the hydrophilic loop (Supplemental Figure S1). While PpPINA and PpPINB share a sequence identity above 86 %, for PpPINC it is below 65 % compared to the others (Table 1, Supplemental Figure S1). Based on these findings and our phylogenetic reconstruction of PINs from selected plant species (Figure 1), we conclude that PpPINA, PpPINB and PpPINC trace back to a single sequence in the last common ancestor of the Funariaceae and originated from two gene duplication events in the genomic history of Physcomitrella (Lang *et al*., 2018): an older one that separated PpPINC from the common ancestor of PpPINA and PpPINB and a more recent one that separated the latter two. The genomic *PpPINC* sequence is identical between the two ecotypes Gransden and Reute from the 5’UTR to the 3’UTR.

**Figure 1:**
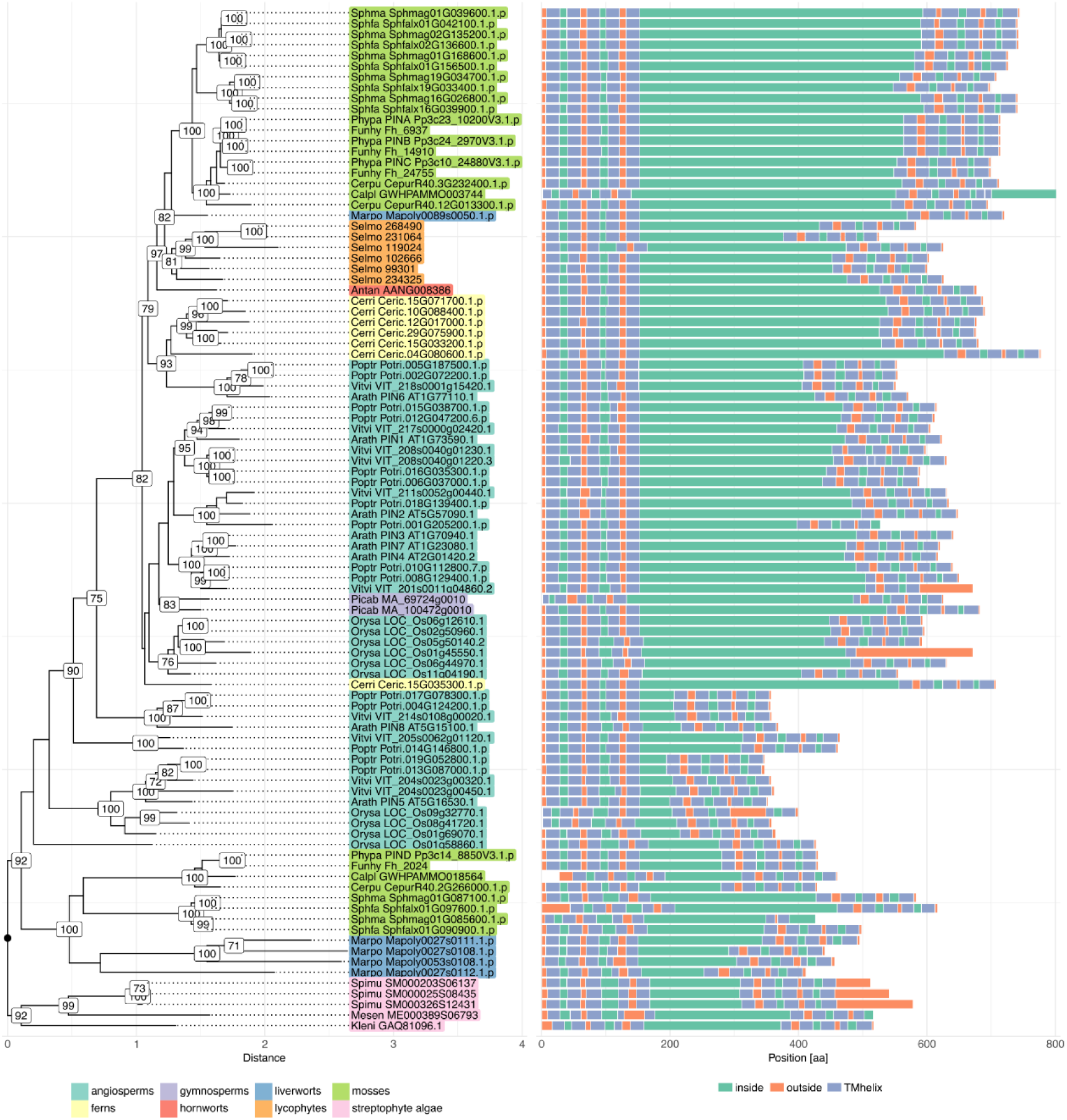
Phylogenetic reconstruction and transmembrane architecture of plant PIN proteins. Left: maximum-likelihood tree of a codon-aware multiple sequence alignment of PIN coding sequences annotated with bootstrap support values (>70 %) and taxonomic units, rooted at the split between streptophyte algae and embryophytes. Right: protein architecture based on DeepTMHMM transmembrane predictions. Species abbreviations: Antan, *Anthoceros angustus*; Arath, *Arabidopsis thaliana*; Calpl, *Calohypnum plumiforme*; Chlre, *Chlamydomonas reinhardtii*; Cerpu, *Ceratodon purpureus*; Cerri, *Ceratopteris richardii*; Funhy, *Funaria hygrometrica*; Kleni, *Klebsormidium nitens*; Marpo, *Marchantia polymorpha*; Mesen, *Mesotaenium endlicherianum*; Orysa, *Oryza sativa*; Phypa, *Physcomitrium patens*; Picab, *Picea abies*; Poptr, *Populus trichocarpa*; Selmo, *Selaginella moellendorffii*; Sphfa, *Sphagnum fallax*; Sphma, *Sphagnum magellanicum*; Spimu, *Spirogloea muscicola*; Vitvi, *Vitis vinifera*.

**Table 1:** Identity, similarity and length of Physcomitrella PpPIN protein motifs

|  | A vs B | A vs C | B vs C | PpPINA | PpPINB | PpPINC |
| --- | --- | --- | --- | --- | --- | --- |
|  | Identity; Similarity of AA in % |  |  | Length AA |  |  |
| TM N' | 96.1;98.7 | 87.7;93.5 | 88.4;93.5 | 155 | 155 | 155 |
| HL | 86.3;91.4 | 64.7;76.9 | 64.4;76.8 | 409 | 409 | 394 |
| TM C' | 95.3;98.7 | 80.0;88.0 | 77.3;86.7 | 150 | 150 | 150 |

A = PpPINA, B = PpPINB, C= PpPINC, TM N' = N-terminal transmembrane region, HL = Hydrophilic loop, TM C'=C-terminal transmembrane region.

Compared to the canonical Arabidopsis PINs (AtPIN1,2,3,4,7), the hydrophilic loop of Physcomitrella PINs are between 51 and 97 AA longer, while there is strong conservation in the transmembrane regions (Supplemental Figure S1).

### Stage-specific expression of *PpPINC*

We compared *PpPIN* expression in publicly available data (PEATmoss database; Fernandez-Pozo *et al*., 2020) for the two Physcomitrella ecotypes Gransden and Reute in two or three datasets, respectively (Hiss *et al*., 2014; Ortiz-Ramírez *et al*., 2016; Perroud *et al*., 2018). As reported in Bennett *et al*. (2014b), *PpPINA* is the highest expressed gene of the three, followed by *PpPINB* with lower, but similar expression. Consistent across the data sets, *PpPINC* is the lowest expressed of the three. While there are some differences in the expression in single tissues in the different data sets, the overall expression of *PIN* genes in both Physcomitrella ecotypes is very similar. In protonema, the expression of all three *PIN*s is the lowest, while the highest expression of *PpPINA* and *PpPINB* can be found in gametophores and developing sporophytes. For *PpPINC,* the expression in vegetative (gametophytic) tissues is very low, while there is a dynamic expression during sporophyte development in both ecotypes, which is nonetheless lower compared to the other two PINs in the same sporophytic tissues (Supplemental Figure S2).

For a closer look, we created a moss line expressing citrine (*pinCPromCit*) under the influence of the native *PpPINC* promoter region (2.1 kb upstream CDS start). This construct was targeted to the carbonic anhydrase-citrine tagged locus created by Wiedemann *et al*. (2018), because of the high expression of the carbonic anhydrase gene and the ease of selection. We screened for altered citrine signals in transformed plants compared to the ever-present citrine expression of the parental line and thus recovered a line with targeted integration of the *pinCPromCit* construct (Supplemental Figure S3). We also created knock-in reporter lines, targeting the hydrophilic loop of PpPINC, similar to localization approaches used in Arabidopsis. However, we found that the transcript of these knock-in lines stopped before the reporter sequence started, resulting in transgenics with no visible reporter signal and no perturbed phenotypes. We therefore dismissed this approach and concentrated on the native promoter strategy. Under standard growth conditions we did not observe *PpPINC*-driven citrine fluorescence, neither in protonema, rhizoids, stems, nor in adult phyllids (Figure 2a). In contrast, citrine fluorescence was clearly visible in the apices of young phyllids. From here, fluorescence proceeds towards the base until maturation of the phyllid. Intriguingly, *PpPINC* expression was highest in parts of the midrib (costa) and in phyllid margins (Figure 2b, c).

**Figure 2:**
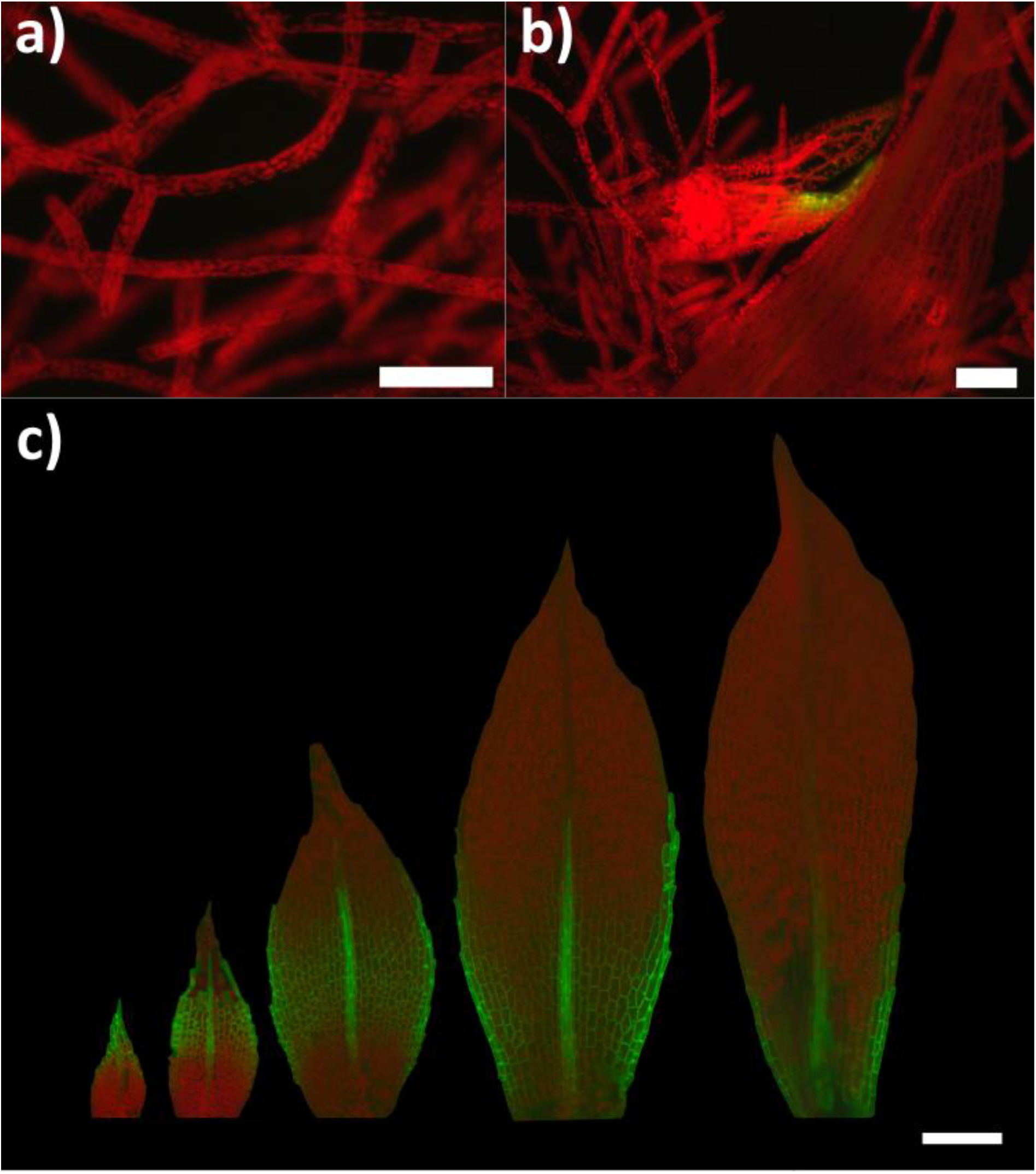
*PpPINC* is expressed in developing phyllids of Physcomitrella. Fluorescence microscopy to visualize citrine expression in a *PpPINC* promoter line (green). Red marks autofluorescence of chlorophyll: a) protonema (bar = 100 µm) b) budding gametophore (bar = 100 µm) c) the youngest phyllids of a moss gametophore (bar = 200 µm).

In addition, citrine expression was detectable in developing gametangia. In mature antheridia close to releasing their sperm cells (antheridium stage 9 in Landberg *et al*., 2013), the signal was found in the foot cells separating the antheridial body from the gametophore apex (Figure 3a). In archegonia, a citrine signal was visible only after the neck canal had opened, surrounding the transition zone between canal and archegonial body above the egg cell (Figure 3b).

**Figure 3:**
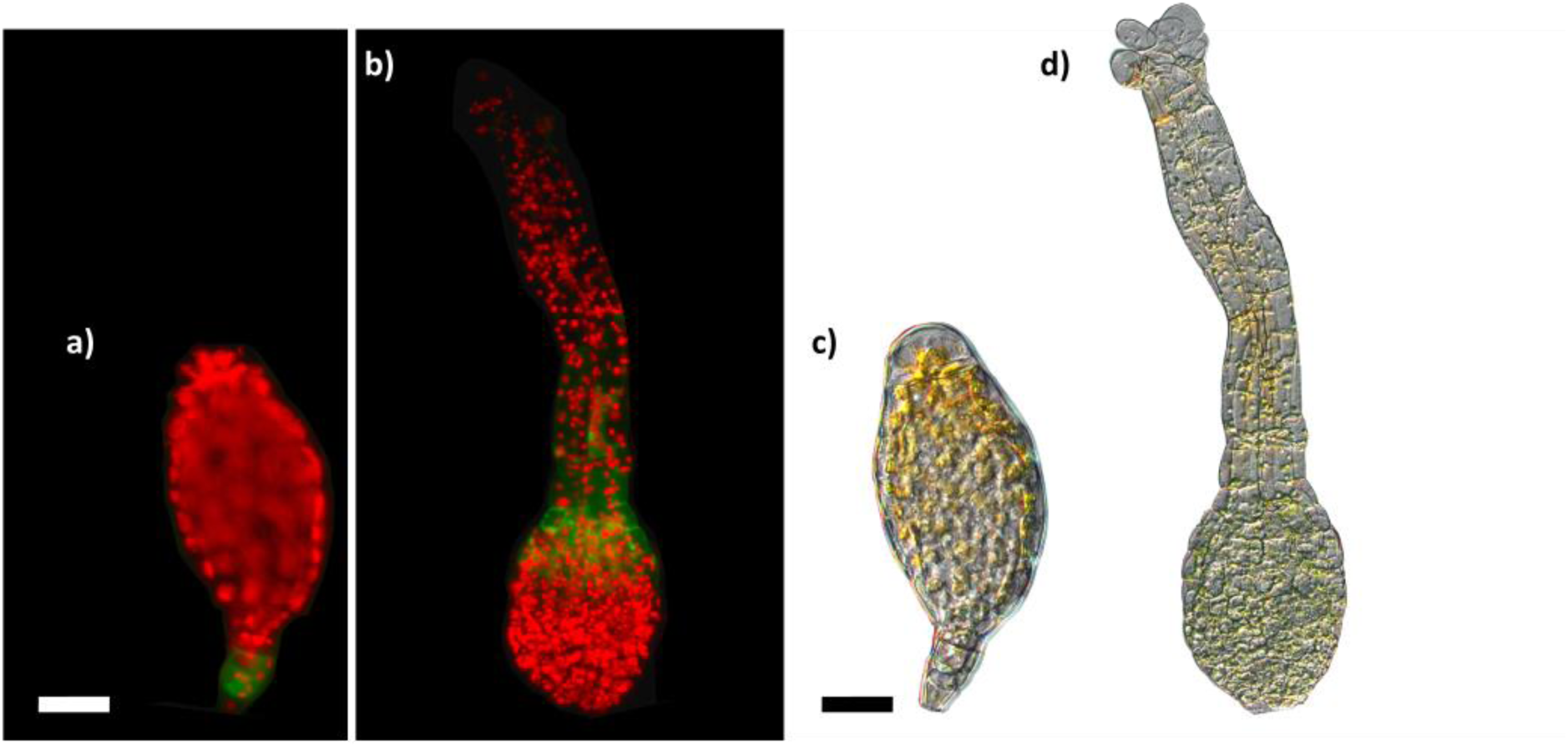
In Physcomitrella, *PpPINC* is dynamically expressed in reproductive organs. Fluorescence-microscopy to visualize citrine expression in a *PpPINC* promoter line (green). Red marks autofluorescence of chlorophyll: a) and c) mature antheridium shortly before sperm cells are released (scale bar = 25 µm). Sperm cells can be seen inside the antheridium in c). b) and d) mature and opened archegonium (scale bar = 50 µm).

### Unaltered vegetative growth in *pin*C mutants

To understand the role of *PpPINC* in the Physcomitrella life cycle, we created targeted knockout mutants via homologous recombination in the background of a fertile Physcomitrella WT. Out of 12 targeted mutant lines devoid of *PpPINC* expression, three independent, haploid mutant lines (*pin*C#10t, *pin*C#29, *pin*C#69) with single integration of the knockout construct in the genome were chosen for further analysis (Supplemental Figure S4). Ploidy was assessed with flow cytometry (Heck *et al.,* 2021) and transgene integration numbers were assessed with the qPCR protocol established by Noy-Malka *et al*. (2014). On solid medium we could not detect obvious phenotypic differences to WT regarding stem length, leaf length and leaf width as well as overall growth (Supplemental Figure S5).

### Altered sperm motility and phenotype in *pin*C mutants

When WT and *pinC* mutants were grown in gametangia-inducing conditions, i.e. 15°C and short day (Hohe *et al*., 2002), male and female gametangia developed without any observable differences. Nevertheless, we detected an increased motility of *pin*C mutant sperm cells (spermatozoids). In WT, 38.9 ± 1 % of spermatozoids are motile 5 minutes after release from the opened antheridium. All three *pin*C mutant lines showed a significantly increased sperm motility (p < 0.00001) of more than 60 % compared to WT (*pin*C#10 = 64.8 ± 2.75 %, *pin*C#29 = 63.8 ± 1.51 %, *pin*C#69 = 62.3 ± 2.32 %) (Figure 4a, b, c). For a deeper analysis we focused on line *pin*C#29. When looking closer at the phenotype of the spermatozoids we observed that a majority of WT spermatozoids have coiled flagella (59.8 ± 11.96 % coiled) (Figure 4d, g, h), while in the mutant *pin*C#29 the vast majority of spermatozoids have non-coiled flagella (94.48 ± 5.22 %) (Figure 4e, f). It was shown by Ortiz-Ramírez *et al*. (2017) that calcium concentration in Physcomitrella sperm cells can alter their motility. However, staining with the calcium-sensitive dye Fluo-4 did not reveal differences in calcium concentrations inside the sperms of WT and mutant *pin*C#29 (Supplemental Figure S6).

**Figure 4:**
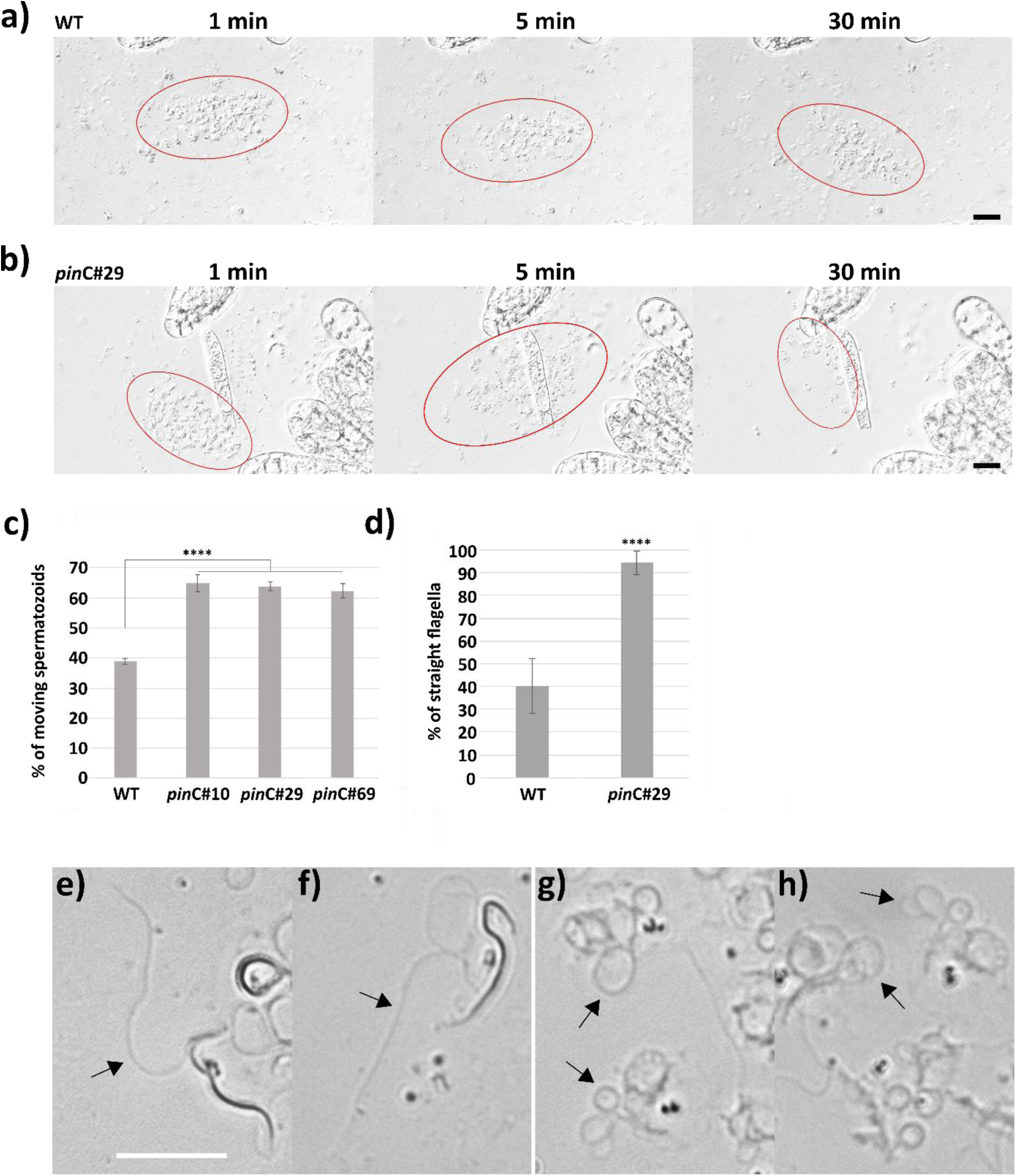
Sperm morphology and motility in Physcomitrella WT and *pin*C mutants. Mutant sperm cells are more motile and have less coiled flagella compared to WT sperm cells. a) WT and b) *pin*C#29 spermatozoids one, five and 30 minutes after being released from a single antheridium, respectively (bar = 20 µm). Areas circled in red highlight spermatozoids after release. c) Percentage of moving spermatozoids after being released from the antheridium (n = 10). d) Percentage of non-coiled flagella compared to coiled flagella (WT: n = 7 antheridia, *pin*C#29: n = 9 antheridia). Asterisks in c) and d) = p ≤ 0.0001, one-sided ANOVA. e) and f) *pin*C#29 spermatozoids with non-coiled flagella marked by arrows. g) and h) WT spermatozoids with coiled flagella marked by arrows. e) – h) bar = 10 µm. For better resolution of sperm cells pictures in a, b, e-h are stacked pictures.

### Organ-specific differential gene expression

To identify genes underlying the differences in sperm flagella phenotype, we performed RNAseq analysis on WT and mutant gametangia. We collected mature archegonia and antheridia from WT and the *pinC*#29 mutant, cultivated on the same plate, 20 days after start of sporophyte induction. Using RNAseq protocols designed for single cell transcriptomics for each line and organ, we pooled eight mature gametangia per sample which were collected on three different occasions (three samples per organ and line). With this approach we could use handpicked gametangia at a specific point in their development. Mapping of sequenced samples resulted in alignment rates of 71.8 – 93.8 % with version 3.3 of the Physcomitrella genome (Lang *et al*., 2018), which is in line with results on Physcomitrella phyllid cells (Kubo *et al.,* 2019) and is comparable to alignment rates for the human genome (Dobin *et al.,* 2013). With 150 bp paired-end Illumina platform-based sequencing, we reached read counts between 19 and 39.8 million for the feature coding sequence (CDS) (Supplemental Figure S7). No reads could be mapped to the deleted area of *PpPINC* in mutant *pin*C#29, which confirms the gene knockout (Supplemental Figure S8). Between all mutant and all WT samples, we could not find significant differences in gene expression and no gene that may have been affected by a possible off-target integration of the knockout construct. In contrast, we found a clear separation between male and female gametangia, while the difference between WT and *pin*C mutant was not strong (Figure 5a). When comparing the samples to each other, we found that for the male samples (WT antheridia WM vs. mutant antheridia PM) there were two upregulated (*Pp3c26_6020, Pp3c26_3990*) and nine downregulated genes (*Pp3c9_8920, Pp3c14_8940, Pp3c20_22670, Pp3c1_22810, Pp3c19_15670, Pp3c3_4950, Pp3c3_11110, Pp3c21_8410, Pp3c12_11710*) (p ≤ 0.05, Fold change (FC) > ± 2, Table 2) (Supplemental Table S2a). Three of the downregulated genes are not annotated so far, while the other genes show counts only in a local part of the gene or exhibit obscure gene structures consisting of only one exon or 22 micro exons. Comparing the female samples, we found one upregulated (*Pp3c11_4360*) and two downregulated (*Pp3c7_8820, Pp3c6_26100)* genes (Supplemental Table S2b), while these genes also show expression only in one part of the gene or show unusual gene structures. When comparing samples derived from antheridia against archegonia, we observed a strong upregulation of genes in male gametangia of both lines, but no change in expression of genes related to female development. The ratio of upregulated to downregulated genes is higher than 4. The highest number of upregulated genes was found in the comparison between all male and all female samples with 1920 genes upregulated and 397 downregulated genes (Table 2). The results of all differentially expressed genes (DEGs) are compiled in Supplemental Table S3.

**Figure 5:**
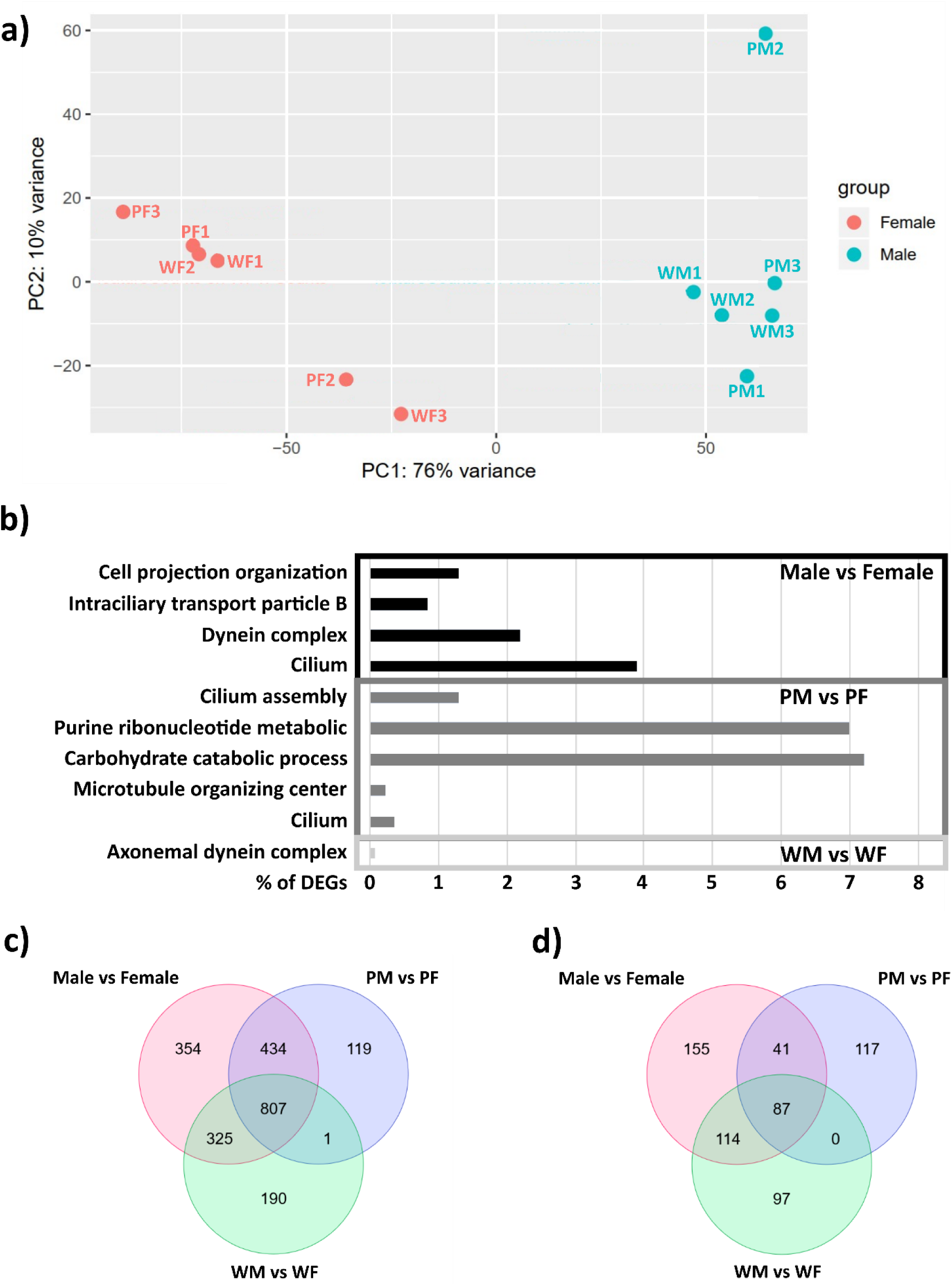
Transcriptomic analysis of Physcomitrella WT and mutant gametangia. a) Variance of all samples. Samples group together into male and female, but there is no clear separation between WT and mutant samples. M = male (antheridia), F = female (archegonia), W = wild type, P = *pin*C#29. b) All enriched gene ontology terms in upregulated DEGs in the comparison of all male against all female samples (male vs. female), all WT male against WT female (WM vs. WF) and mutant male against mutant female samples (PM vs. PF). c) Venn diagram of all upregulated and d) downregulated genes in the comparisons of male and female samples.

**Table 2:** Upregulated and downregulated genes in male (M) and female (F) tissue samples from Physcomitrella WT (W) and pinC#29 (P).

|  | Upregulated genes | Downregulated genes |
| --- | --- | --- |
| **PM vs. WM** | 2 | 9 |
| **PF vs. WF** | 1 | 2 |
| ***pinC#29* vs. WT** | 0 | 0 |
| **PM vs. PF** | 1361 | 245 |
| **WM vs. WF** | 1323 | 298 |
| **Male vs. Female** | 1920 | 397 |

We analysed the gene ontology (GO) terms of the DEGs for accumulation of specific terms for all experiments (p-value cut off 0.01). Enriched GO terms occurred in the comparisons between sexes, especially in upregulated genes (Figure 5b). For the comparison of WT antheridia against WT archegonia (WM vs. WF) we found a weak enrichment of the GO term axonemal dynein complex (0.08 %). In the experiment PM vs. PF (mutant antheridia vs. mutant archegonia) we found that five GO terms were most prominent, three are associated with spermatogenesis and one each with energy consumption and DNA synthesis (Cellular Components: cilium = 0.36 %, microtubule organizing center = 0.23 %; Biological Process: carbohydrate catabolic process = 7.2 %, cilium assembly = 1.3 %, purine ribonucleotide metabolic process = 7 %). The same is true for the comparison of all antheridia samples with all archegonia samples, where four spermatogenesis-related GO terms were overrepresented (Cellular Components: cilium = 3.9 %, dynein complex = 2.2 %, intraciliary transport particle B = 0.84 %; Biological Process: cell projection organization = 1.3 %) (Supplemental Table S1).

In consequence, we detected more spermatogenesis-related DEGs in the comparison between mutant antheridia and archegonia than in the comparison of WT gametangia. Most of the upregulated DEGs found in the comparison of the sexes were shared between WT and mutant (Figure 5c). The number of DEGs found exclusively in the mutant was similar between upregulated and downregulated genes (119 / 117). The WT shared the least downregulated genes with the mutant, while contributing more downregulated DEGs to the comparison between male and female (Figure 5d). We identified some single DEGs, which had been reported to play roles in flagella formation or auxin homeostasis. The coiled coil-like protein *Ppccdc39* (Meyberg *et al*., 2020) was upregulated almost 3-fold (FC) in male samples compared to female samples (FC = 2.98). While it is also significantly upregulated in the comparison of WT male against female gametangia (FC = 2.4, p = 0.0005), the fold change was even larger in the mutant samples, but due to high variation in the male samples not statistically significant. The arl13b homologue *Pp3c1_40600*, which is involved in flagella stability, was significantly upregulated in the comparison of male against female gametangia (FC = 3.44, p = 0.025), which is also upregulated in the Reute ecotype compared to Gransden (Meyberg *et al*., 2020). In all comparisons of male against female tissues the arabinogalactan 31 homologue *Pp3c5_9210*, found to be active during spermatogenesis (Meyberg *et al*., 2020), was upregulated with a higher fold change in PM vs. PF (FC = 7.24, p = 2.06E-10) than in WM vs WF (FC = 4.62, p = 1.20E-07) (male vs. female FC = 5.34, p = 1.61E-08). Furthermore, the embryogenesis-related transcription factor *PpBELL2* (Horst *et al*., 2016) was significantly downregulated in all male samples compared to female samples (PM vs. PF: FC = - 7.53, p = 8.17E-09; WM vs. WF: FC = −3.07, p = 0.013, male vs. female: FC = −3.85, p = 0.0029). The two PHD clade IIa genes *PpMS1A* and *PpMS1B* are expressed significantly higher in male tissues compared to female (*PpMS1A*: male vs. female FC = 2.35, p = 0.029, *PpMS1B*: male vs. female FC = 2.81, p = 0.036, WM vs WF FC = 2.04, p = 0.035), which is in accordance with Landberg *et al*. (2022).

While not being differentially expressed, we observed expression of all six *PpTAR* genes in antheridia and archegonia. We could also find higher expression levels of *PpYUCB* and *PpYUCF* in antheridia and archegonia compared to the other *PpYUC* genes, while *PpYUCD* showed a very low, but significantly upregulated expression in archegonia (PM vs. PF FC = −4.69, p = 0.02, Male vs. Female FC = −4.65, p = 0.0008). This indicates active auxin synthesis in mature gametangia. While our transcriptomic data reveal trends in general gene expression, we could not identify a single DEG that might account for the difference in sperm flagella phenotype in the comparison of WT and mutant, but found more spermatogenesis-related DEGs in the *pin*C mutant.

### Altered fertility and abortion rate

We did not observe any differences in the morphology of mature spore capsules or spore germination rates between WT and mutant (Supplemental Figure S9). In contrast, in addition to the higher motility of mutant spermatozoids, we found a significantly higher fertility rate in all three mutant lines (Figure 6). Low fertility rates of the Gransden ecotype have been reported (Perroud *et al*., 2011; Hiss *et al*., 2017; Meyberg *et al*., 2020), and in our experiments 5.5 ± 0.4 % of all WT gametophores produced a sporophyte. In contrast, all *pin*C mutants developed significantly more sporophytes, with 39.8 ± 1.6 % for *pin*C#10, 63.2 ± 4.5 % for *pin*C#29 and 14.5 ± 0.6 % for *pin*C#69.

**Figure 6:**
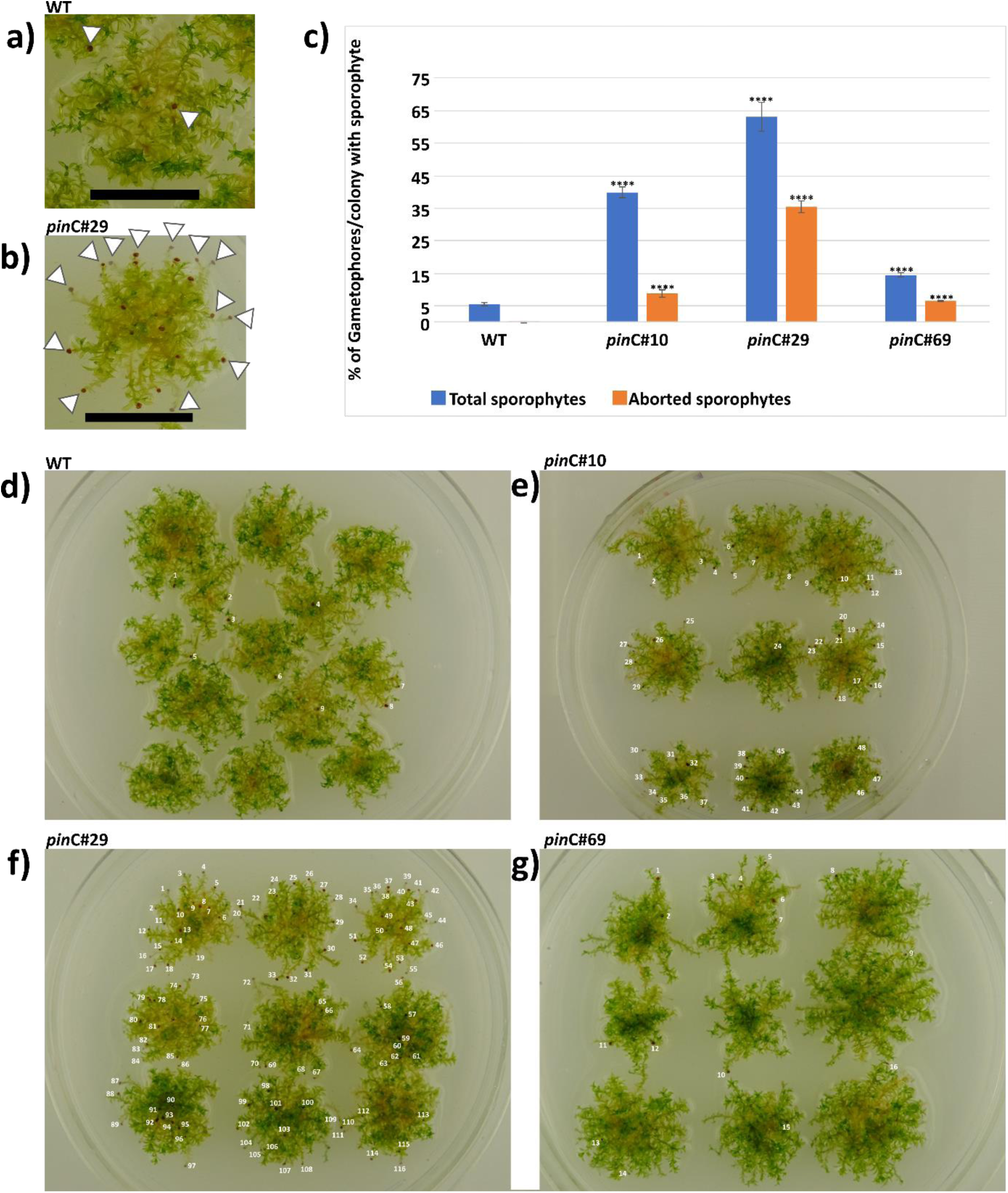
Spore capsule numbers. Physcomitrella spore capsules in a) WT and b) *pin*C#29 mutant, white arrows mark sporophytes (for visibility reasons, not all sporophytes in b) are marked while all visible sporophytes are marked in a)). bar = 1cm. c) Percentage of gametophores with a sporophyte per colony. Total number of sporophytes (adult + aborted) in blue, only aborted sporophytes in orange. Asterisks = p<0.00005 (Student’s t-test) of mutant lines against WT, n (colonies) = WT (41), *pin*C#10 (30), *pin*C#29 (42), *pin*C#69 (32) d) WT plate with 9 sporophytes counted. e) *pin*C#10 plate with 48 sporophytes. f) *pin*C#29 plate with 116 sporophytes. g) *pin*C#69 with 16 sporophytes.

In addition to increased fertility, we detected an increased abortion rate of sporophytes in the mutants. For this we examined every gametophore apex of a colony and counted sporophytes, empty vaginulae and apices with no gametangia. Abortion rate was determined for each colony by calculating the percentage of aborted sporophytes per gametophore per colony. Subsequently, we calculated the abortion rates in WT and mutants to be WT = 0.1 ± 0.03 %, *pin*C#10 = 8.7 ± 1.3 %, *pin*C#29 = 35.4 ± 1.7 %, and *pin*C#69 = 6.5 ± 0.2 % (Supplemental Table S4). The aborted sporophytes were no longer attached to the maternal tissue (vaginula). Abortion happened around two weeks after fertilization of the egg cell, when the sporogenic tissue starts to expand (Stage S3 in Ortiz-Ramírez *et al*., 2016; day 12-16 in Lopez-Obando *et al*., 2022).

In the early embryo, *PpPINC* was active in the lower half, excluding the basal tip cells, as well as in the walls of the maternal tissue that surrounds the young embryo (epigonium) (Figure 7a). Development of the sporophyte foot and the maternal tissue are highly synchronized. When the embryo has doubled in size and the foot is secured in the now fully developed vaginula (Figure 7b), the seta forms and rips apart the surrounding tissue of the epigonium, while the later developing spore capsule also starts to separate (Figure 7c), splitting the epigonium into the calyptra at the apex and the vaginula at the basis of the premeiotic sporophyte. The expression of *PpPINC* slowly declines in the foot of the embryo (Figure 7a-c), while it increases in the maternal tissue during sporophyte foot growth, forming a distinct ring structure at the border of maternal and sporophytic tissue (Figure 7b). After the growth spurt, *PpPINC* expression can be found only in the apophysis (region between seta and premeiotic tissue), while excluding the stomata cells (Figure 7d). We found that Physcomitrella develops a true-type vaginula, where the foot of the sporophyte does not penetrate the gametophore tissue under the vaginula (Figure 7e, f). Sporophyte development is polar in Physcomitrella, where the foot of the sporophyte develops faster than the seta or premeiotic spore capsule. The basis of the foot was not in direct contact with the maternal cells, which is in line with an earlier report (Regmi *et al*., 2017), while the vaginula tightened at its apical border to the sporophyte, which is visible as a reddish-brown coloured ring after the emergence of the premeiotic sporophyte (Figure 7g, i). In the mature sporophyte, no *PpPINC* expression was detectable.

**Figure 7:**
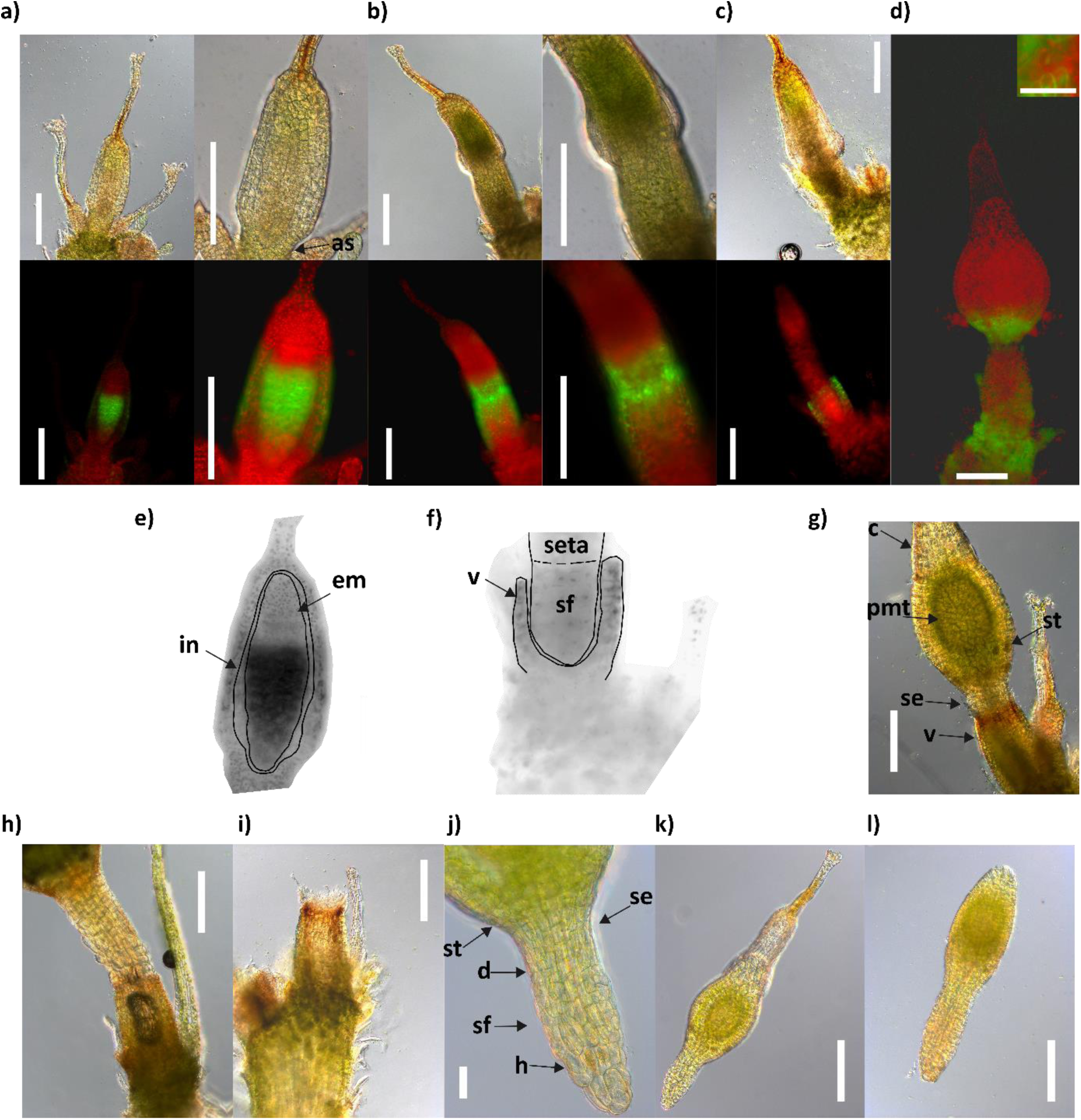
Sporophyte development in Physcomitrella. a)-d) Bright-field and fluorescent microscopy pictures of Physcomitrella PpPINC reporter line. a) Embryo (probably 128 cell stadium) inside the epigonium, as = archegonial stalk. b) Embryo has doubled in size, vaginula and sporophyte foot are fully developed. c) Growth phase of seta, epigonium is about to rupture. d) Premeiotic sporophyte around 14 days post fertilization. The fluorescent signal is concentrated in the apophysis excluding stomata cells (smaller picture, bar = 25 µm). e) and f) = Negative grayscale cut out of a) = e) and c) = f), e) em = embryo, and in = inner wall epigonium are outlined in black. f) Fully developed vaginula (v) with sporophyte foot (sf) and growing seta are indicated by black lines. g) Premeiotic sporophyte, after the epigonium has split into the calyptra (c) and vaginula (v), pmt = premeiotic tissue, se = seta, st = stomata. h) Premeiotic sporophyte slips out of vaginula. i) Empty vaginula after abortion. j) Lower half of an aborted premeiotic sporophyte, d = depression, h = haustorium cells, se = seta, sf = sporophyte foot, st = stomata). k) Aborted premeiotic sporophyte covered by calyptra. h) – k) = *pin*C#29 mutant. l) Aborted premeiotic sporophyte from WT, not attached to calyptra. Bars in a), c), d), g), h), i), k), and l) = 200 µm. Bar in j) = 50 µm.

Aborted sporophytes in WT and mutants were in the premeiotic phase after the separation of the epigonium (comparable to stage shown in Figure 7g), and no longer attached to the vaginula. No visible defects could be detected, apart from the separation from the gametophore. On one occasion we could observe a sporophyte slipping out of the cavity of the vaginula, not losing contact as the basis of the foot was stuck in the tighter apical end (Figure 7h). Empty vaginulae clearly showed the coloured ring formation at the apical opening (Figure 7i). The aborted sporophytes had fully developed foot structures, with haustorium cells at the basis, seta, stomata cells and a depression between seta and haustorium cells, which we believe results from the securement of the sporophyte by the coloured ring formation (Figure 7j). We could find no morphological differences in the aborted sporophytes between mutants and WT, except for an increase in the number and percentage of abortions in the *pin*C mutants. The calyptra could be removed without resistance, some aborted sporophytes had already lost them (Figure 7k, l).

## Discussion

Auxin plays crucial roles in plant development (Santner & Estelle, 2009; Weijers & Wagner, 2016), including Physcomitrella development (Decker *et al*., 2006; Menand *et al*., 2007; Fujita *et al*., 2008; Coudert *et al*., 2017; Nemec-Venza *et al*., 2022) and biotechnology (Ruiz-Molina *et al*., 2022). Although PIN proteins are central in auxin transport and action (Adamowski & Friml, 2015; Sauer & Kleine-Vehn, 2019; Ung *et al*., 2022), only two of the three canonical Physcomitrella PIN proteins have been fully characterized; the similar and relatively highly expressed PpPINA and PpPINB. In contrast, little was known about the function of the more divergent and less expressed PpPINC. Here, we strived to close this knowledge gap.

Our data on the expression of *PpPIN*C in young Physcomitrella phyllids is in line with an earlier report (Viaene *et al*., 2014), as well as the absence of the signal in stem, rhizoids and older phyllids. It was known that the single knockout of *PpPINA* or *PpPINB* has no visible effect on the morphology of the gametophore or phyllids (Bennett *et al*., 2014b), and this is also true for *PpPINC*, as we have shown here. However, a double knockout of *PpPINA* and *PpPINB* lead to elongated phyllids and stems, similar to treatments with exogenous auxin or auxin transport inhibitors (Bennett *et al*., 2014b). This means that while *PpPINA* or *PpPINB* together with *PpPINC* can rescue the knockout of either *A* or *B*, *PpPINC* alone cannot replace the function of both proteins in a double knockout line.

The publicly available expression data for canonical PIN proteins in the two Physcomitrella ecotypes Gransden and Reute is not always comparable, as they are different experimental data sets describing specific tissue stages. However, all three canonical PINs have a similar expression pattern regarding tissue and ecotype. All three PINs are expressed in gametophores and sporophytes, with PpPINA and PpPINB being more highly expressed than PpPINC. The expression of all three PINs in vegetative (gametophytic) tissue has been reported (Viaene *et al*., 2014), as well as the important role of PpPINA and PpPINB in sporophyte formation (Bennett *et al*., 2014b). We could not find any differences between the expression of *PpPINC* in the Gransden and Reute data sets. Further, we did not observe any vegetative phenotype alteration in *PpPINC* knockout mutants. While Viaene *et al*. (2014) showed expression of all three canonical PIN proteins in Physcomitrella protonema apical cells, we could not find such expression. This might be due to differences in targeting, reporter construct or culture conditions. In contrast, we could identify PpPINC as an important regulator of spermatogenesis and sporophyte retention. We therefore conclude that PpPINC is functionally active only in gametangia and sporophytic tissues, contrary to the other two canonical Physcomitrella PIN proteins. Consequently, the *pin*AB double mutants are affected in gametophytic as well as sporophytic development (Bennett *et al*., 2014b), while this is not the case in the *pin*C mutants analysed here.

It has been reported that the low fertility of the Physcomitrella ecotype Gransden is based on reduced male fertility (Perroud *et al*., 2011). This is partially caused by coiled flagella of spermatozoids, which results in low sperm motility (Meyberg *et al*., 2020). We confirm this phenotype for the Gransden ecotype, with a high percentage of coiled flagella and a motility of under 50 %, resulting in a very low sporophyte production rate. In contrast, the *pin*C mutants, which we generated in the Gransden background, resemble the Reute ecotype in sperm morphology and fertility rate, as it has no coiled flagella, a high sperm motility and a high sporophyte rate (Hiss *et al*., 2017; Meyberg *et al*., 2020). While the fertility rate of the three *pin*C mutant lines varied, it was significantly higher in all three lines than in WT. There are many biological factors at play which could influence fertilization success on the female side. However, one essential factor for successful fertilization is the ability of sperm cells to swim to an egg cell. Therefore, it is evident that higher sperm motility leads to higher fertilization rates, as we observed here in the *pin*C mutants. The Gransden ecotype was introduced by Engel (1968) as a laboratory strain from one single spore from the UK, and has been propagated mostly vegetatively in laboratories around the world since, while the Reute ecotype was introduced relatively recently as a collection from Germany (Hiss *et al*., 2017). Compared to Reute, Gransden accumulated somatic, epigenetic mutations, probably leading to faults in spermatogenesis (Haas *et al*., 2020).

Here, we found more spermatogenesis-related DEGs in mature antheridia of a *pin*C knockout mutant compared to WT (WT: axonemal dynein complex = 0.08 %, *pin*C#29: cilium = 0.36 %, microtubule organizing center = 0.23 % cilium assembly = 1.3 %), but we could not find changes regarding female or egg-cell related differences in expression in mature mutant and WT archegonia. Activity of a *TAR* gene and *PpPINA* in the apical cells of the mature antheridium has been reported (Landberg *et al*., 2013; Landberg *et al*., 2020), while there are no reports of phenotypically different gametangia in *PpPINA* mutants. Here, we also found evidence for expression of auxin synthesis-genes in mature gametangia. With *PpPINC* being expressed in the foot of the mature antheridium, contrary to *PpPINA* in the apical tip cell, it seems to be more important for spermatogenesis to control auxin homeostasis at the bottom of the antheridium than at the tip. Whether this mode of action is part of a polar auxin homeostasis in a moss organ controlled by PIN proteins needs further clarification.

Gaining the ability to move the flagellum is one of the final steps of spermatogenesis before spermatozoids are released and activated. In mammals, this ability is gained in the epididymis, and is controlled by different external and internal factors (Pereira *et al*., 2017; O’Flaherty, 2019; Björkgren & Sipilä, 2019). Due to the complexity of the process and a large number of influences, sperm populations are not homogenous, but vary regarding phenotype, motility or activity (Gómez Montoto *et al*., 2011; Genau *et al*., 2021; Martins-Bessa *et al*., 2021). A key role in metazoan and Physcomitrella spermatogenesis is played by the evolutionary conserved DNA Topoisomerase 1α, which facilitates chromatin condensation towards the compact sperm head (Gu *et al*., 2022). Activation of the flagella of released spermatozoids depends on changes in pH, calcium concentration, or the presence of a chemoattractant released by the egg cell (Nakajima, 2005; Suarez, 2008; Morita *et al*., 2009). While mammalian spermatozoids are transported through the epididymis during maturation, spermatogenesis in the moss antheridium is stationary. This increases pressure on the exact spatio-temporal expression of spermatogenesis-related genes. As in mammals, Physcomitrella releases heterogeneous sperm populations from one antheridium. This was true for WT where motility (∼40 %) and non-coiled flagella (∼40 %) seem to fit nicely, whereas in the *pin*C#29 mutant over 90 % of all spermatozoids were straight, with an overall motility of 60 %. While the phenotype of the spermatozoids changed drastically in the mutant, motility did not increase at the same rate. As we have shown, *PpPINC* is expressed only shortly before sperm cells are released, reducing the time it can influence spermatogenesis to a short window. Therefore, alterations in duration and strength of expression, which are likely to occur after somatic mutations, could explain the differences in sperm morphology between the two ecotypes Gransden and Reute. The *PpPINC* gDNA sequence between the Gransden ecotype (v3.3 Phytozome genome ID:318) and Reute (SRX1528135; Hiss *et al*., 2017) is identical. Given that the mutant antheridia exhibit a higher portion of spermatogenesis-related DEGs compared to WT, together with an overall increased motility and fertility, we suggest that PpPINC acts in repression of spermatogenesis. The repression of spermatogenesis at the end of the whole process could be a molecular signal for sperm release or activation of flagella. The difference in expression between both ecotypes would be an earlier repression in Gransden, halting spermatogenesis when most of the flagella are coiled and not yet ready for release, while the signal in Reute comes later, when spermatogenesis has progressed to a majority of non-coiled sperm flagella. The unknown activating signal of *PpPINC* expression could therefore be the culprit responsible for reduced male fertility in the Gransden ecotype. The identification of such regulators is in its infancy, with WAV3 in the regulation of Arabidopsis PIN2 being the first (Konstantinova *et al*., 2022).

Male sexuality is reduced in many bryophytes, with a female-biased sex ratio (Cameron & Wyatt, 1990; Stark *et al*., 2010; Pépin *et al*., 2013; Bisang *et al*., 2015; de Jong *et al*., 2018), or size of the plants, with the occurrence of dwarf males (nannandry), which are unique in bryophytes among land plants (Pedersen *et al*., 2006; Rosengren & Cronberg, 2014; Rosengren *et al*., 2016; Lang *et al*., 2021). Dwarf males grow on the leaves of female plants (Pichonet & Gradstein, 2012; Rosengren & Cronberg, 2014; Rosengren *et al*., 2016; Lang *et al*., 2021) and increase fertilization success (Hedenäs & Bisang 2012; Rosengren & Cronberg, 2014), in the absence of a female, male spores develop normally. In *Macromitrium japonicum*, dwarf males grew in culture on medium containing auxin, but developed normally on auxin-free medium (Une, 1985). Dioecious mosses grow mostly vegetatively and sporophyte production can be rare due to absence of a sexual partner, while monoecious mosses produce sporophytes more frequently, as the chances for fertilization are higher (Haig, 2016). However, self-fertilization leads to homozygous spores, while self-produced sperm are rarely outcompeted by non-self sperm (Taylor *et al*., 2007; Rosengren *et al*., 2016). Reducing male fertility in a monoecious moss increases the chance for outcrossing and could therefore be an internal mechanism, controlling the need to refresh genetic material (McDaniel *et al*., 2010; Haig, 2016; Szövényi *et al*., 2017). In a monoecious moss like Physcomitrella with a very short life cycle (3-6 months) (Cove, 2005), pressure on mutations regarding the sexual life cycle is strong, as changes in fertility would be lethal (Haig, 2016). Cultivation in vegetative culture in laboratories around the world (Haas *et al*., 2020) could reduce this pressure on fertility, increasing the risk for severe mutations in the sexual signalling cascade (Meyberg *et al*., 2020; Haas *et al*., 2020). As male gametes need more energy and are more complex to build than female gametes (Rydgren & Økland, 2003; Stark *et al*., 2000, 2009; Horsley *et al*., 2011; Santos *et al*., 2022), risk for mutations is higher. This cost calculation would also favour intentionally reducing male rather than female fertility, as more energy is required to constantly produce sperm cells rather than egg cells, which are waiting for fertilization, ending the gamete production cycle and starting the growth of propagules.

In Arabidopsis, the expression of PIN proteins plays an important role during pollen development, together with anther-specific expression of YUCCA genes (Cecchetti *et al*., 2008; Dal Bosco *et al*., 2012). PIN8 locates to the ER and regulates auxin homeostasis with a rate-limiting activity during pollen grain development and pollen tube growth. It is functionally active only during male gametophyte development and a knockout of PIN8 leads to misshaped and aborted pollen (Ding *et al*., 2012; Dal Bosco *et al*., 2012). The activity of PIN proteins in male gametophytic tissues is also reported in algae, in *Chara vulgaris* a PIN2-like protein is expressed during spermatogenesis (Żabka *et al*., 2016). Together with our findings it seems plausible that PIN proteins can play important roles during male gametophytic development in all plants. However, the exclusive function of PIN1 in Arabidopsis in the formation of floral organs could not be rescued by the Physcomitrella PpPINA protein expressed under the PIN1 promoter, while it complements the vegetative phenotype of the knockout (Zhang *et al*., 2020a).

The abortion of embryos is a natural process. Unfavourable environmental conditions, genetic mutations, injury of the embryo or of maternal tissue can trigger abortion. In mosses, the normal abortion rate differs among species (Stark & Stephenson, 1983; Stark *et al*., 2009; Rosengren *et al*., 2016; Hedenäs & Bisang, 2019) and seems to be resource-limited (Stark *et al*., 2000). Mosses have to allocate their energy between clonal regeneration and sexual reproduction (Stark *et al*., 2009), and sporophyte survival positively correlates with vegetative growth prior to fertilization (Stark & Stephenson, 1983). In these cases, aborted sporophytes were no longer supported with nutrients and stopped growing inside the vaginula, contrary to the active abortion we observed here in the *pin*C mutant.

The haustorium cells of the sporophyte foot are not pressed against the vaginula tissue but surrounded by a placenta-like space, while both tissues are separated by a diffusion barrier (Uzawa & Higuchi, 2010; Regmi *et al*., 2017). The foot of the sporophyte is wider than the seta with a small depression between seta and foot, while vital sporophytes are tightly attached to the gametophore. We observed a clear polarity during early embryo development favouring growth of the foot, while the upper part of the embryo starts to increase only after the foot is secured in the vaginula. *PpPINC* is active in the maternal tissues which will form the vaginula and the sclerotized ring structure after the sporophyte ruptures the epigonium. Premeiotic sporophytes were aborted after the epigonium had ruptured and the sporophyte foot had to be secured in the vaginula. The significantly increased abortion rate in our *pin*C mutants, combined with the activity of *PpPINC* at the vaginula-seta junction, point to a regulation of sporophyte securement controlled at least partially by *PpPINC*. This sporophyte abortion phenotype was not observed with the *pin*AB double mutants, which have malformed sporophytes that are nevertheless firmly attached to the gametophore (Bennett *et al*., 2014b). Based on our observations, we suggest that the sclerotized, brown ring structure at the vaginula-seta junction has the mechanical function of securing the sporophyte foot. It is well-known that auxin can have a loosening effect on cell walls (Cosgrove, 2021), regulated by complex molecular signalling (Xiong *et al.,* 2021; Varapparambath *et al.,* 2022). Thus, changes in auxin distribution or concentration in the ring structure could lead to a loosening of the ring and subsequently to the loss of the sporophyte.

The polarity of early embryo development we observed here in Physcomitrella as well as the functional significance of the sclerotized ring structure is in line with the findings of Lorch (1909) in the moss family Polytrichaceae. He reported that the sporophyte foot develops first, before the seta subsequently elongates, and as the lumen in the vaginula is not completely filled by the foot, the sclerotized ring structure must secure the sporophyte (Lorch, 1909). As we could not find a name for this ring structure in the literature, we propose to name the reddish-brown ring structure, formed at the junction of vaginula and seta, the Lorch ring.

One important step in the evolution of land plants was the retention of the product of fertilization, the zygote, on the haploid gametophyte (Reski, 2018). This “zygote retention” already occurs in streptophyte algae (Becker & Marin, 2009) and is a prerequisite for the development of the alternation of generations that is characteristic for all land plants (Horst & Reski, 2016). Our data presented here suggest that the structure that mechanically ensures sporophyte retention is the Lorch ring, and that retention and abortion of the sporophyte can be regulated by auxin via PpPINC.

Taken together, the canonical Physcomitrella PINC protein is functional in reproductive tissues only, an important regulator of late spermatogenesis, sperm motility and of retention and active abortion of premeiotic sporophytes. Thus, it may integrate environmental signals with developmental programs to regulate sexual reproduction, at least in moss gametangia and early stages of embryo development.

## Acknowledgements

We gratefully acknowledge Richard Haas for expert technical advice and assistance extracting RNA from gametangia, and Anne Katrin Prowse for proof-reading of the manuscript. We gratefully acknowledge funding by the Deutsche Forschungsgemeinschaft (DFG, German Research Foundation) under Germany’s Excellence Strategy EXC-2189 (CIBSS to RR) and EXC-2193/1 – 390951807 (*liv*MatS to RR).

## Author contributions

VML, CR, ELD and RR planned and designed the research. VML did most of the experimental work. NvG performed sequence search and alignment, phylogenetic reconstruction and transmembrane prediction. Data analysis of transcriptomic sequences was done by VML, guided by CR. MB created the knockout constructs and first knockout plants. OH took pictures of spermatozoids and aborted sporophytes. MH did measurements of stems and leaves, while data analysis for this was done by VML. ELD and RR supervised research. RR acquired funding for this work. VML wrote the manuscript with the help of ELD and RR. All authors read and approved the manuscript.

## Declaration

All authors declare that they have no competing interests.

## Data availability

All RNA-seq samples as well as the DEG experiments created in this study are available via the NCBI GEO project **GSE205257**. All moss lines used are available via the International Moss Stock Center (IMSC; https://www.moss-stock-center.org) with the following accession numbers: WT = 40095, *PinCPromCit =* 40917, *pin*C#10 = 40918, *pin*C#29 = 40919, *pin*C#69 = 40420.

## Supporting Information

**Supplemental Figure S1:**
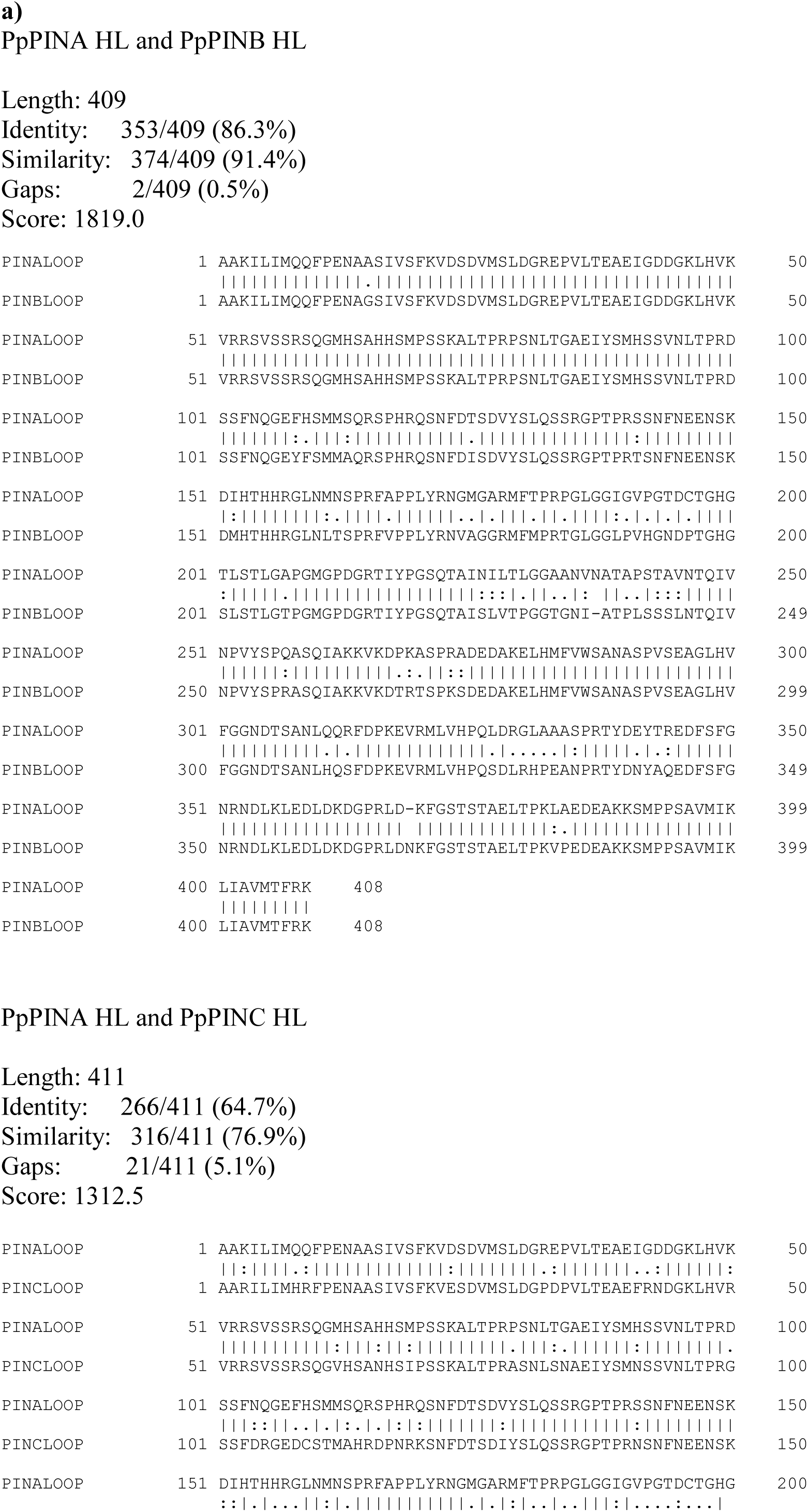
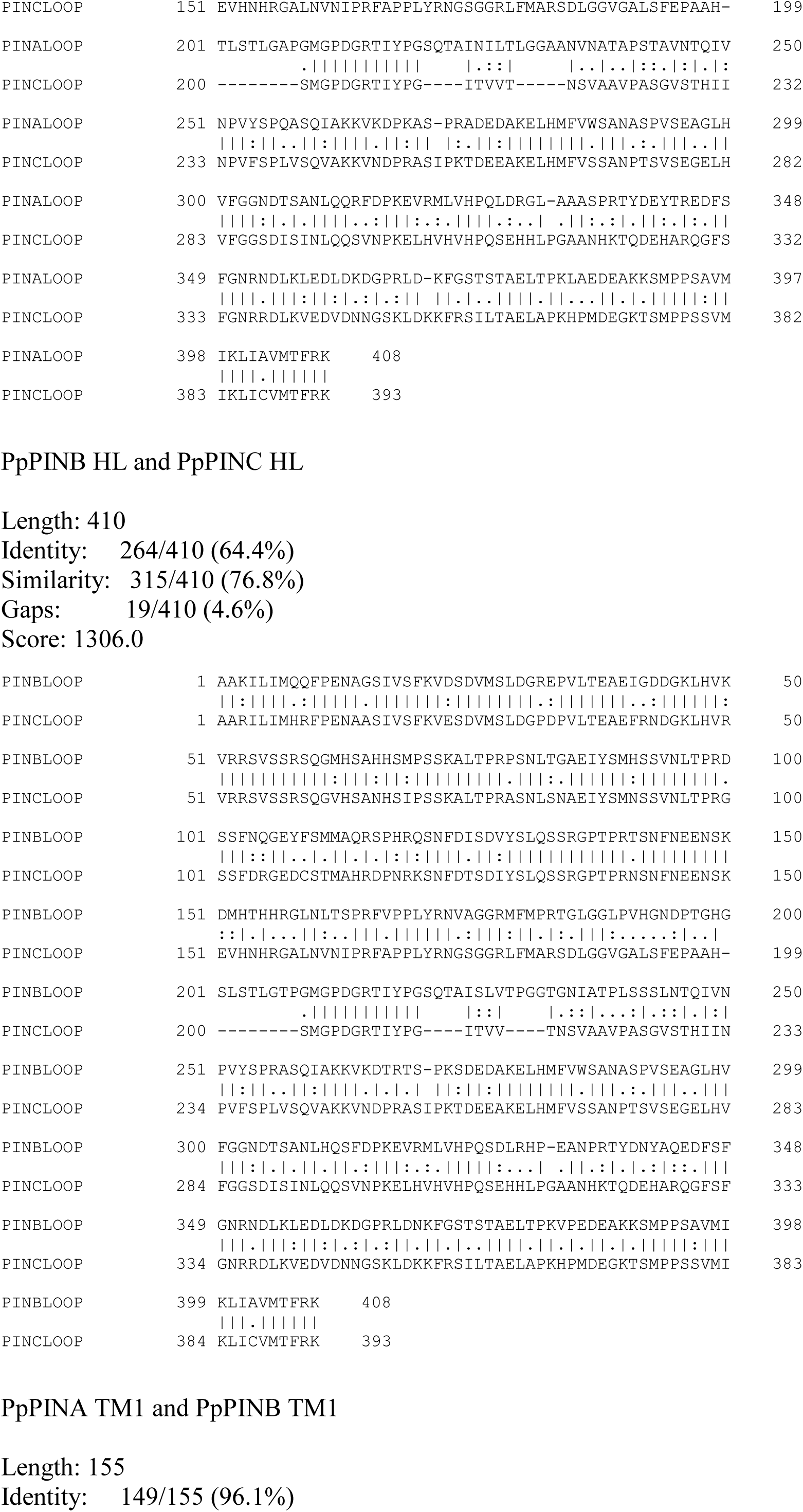
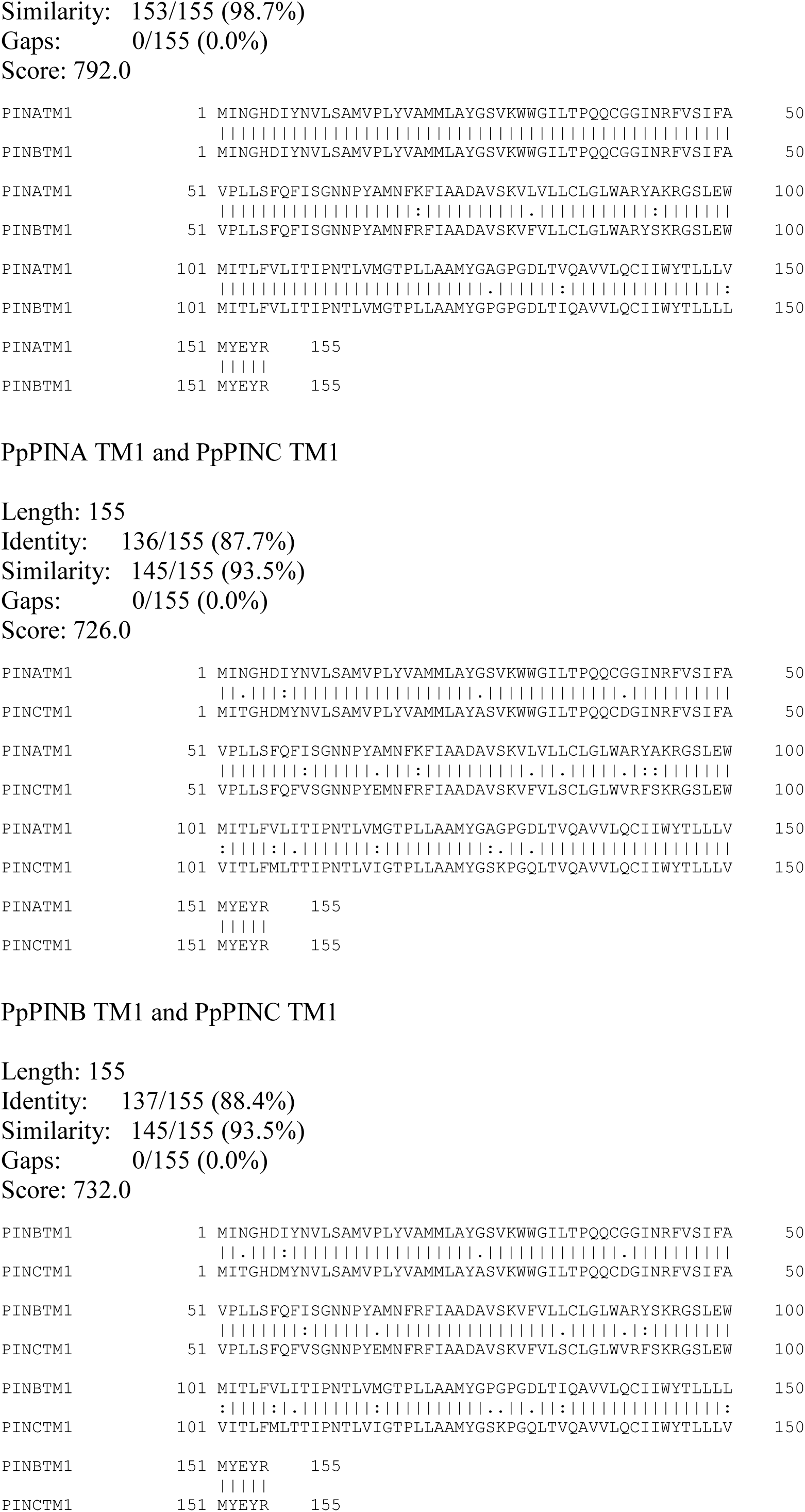
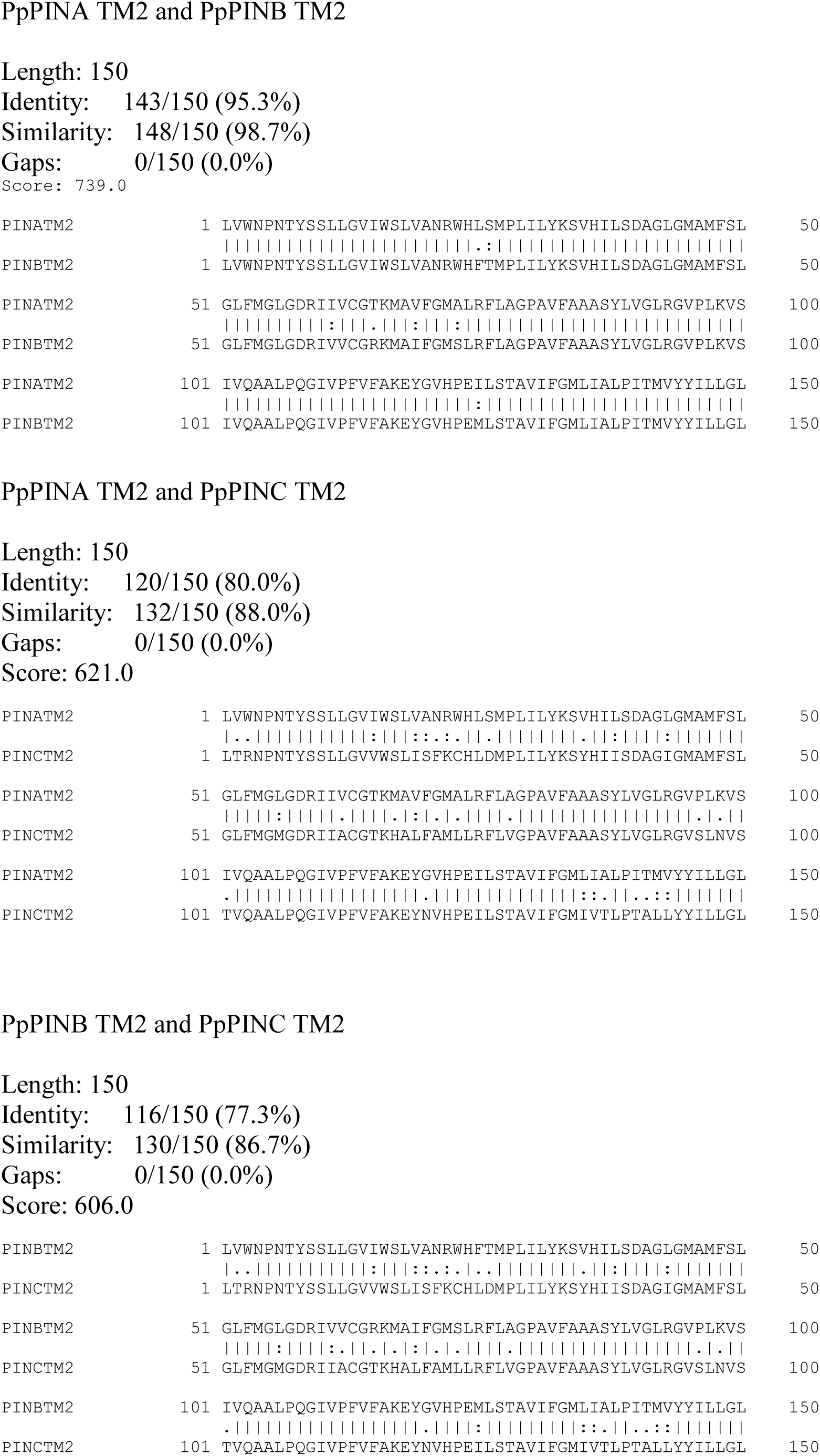
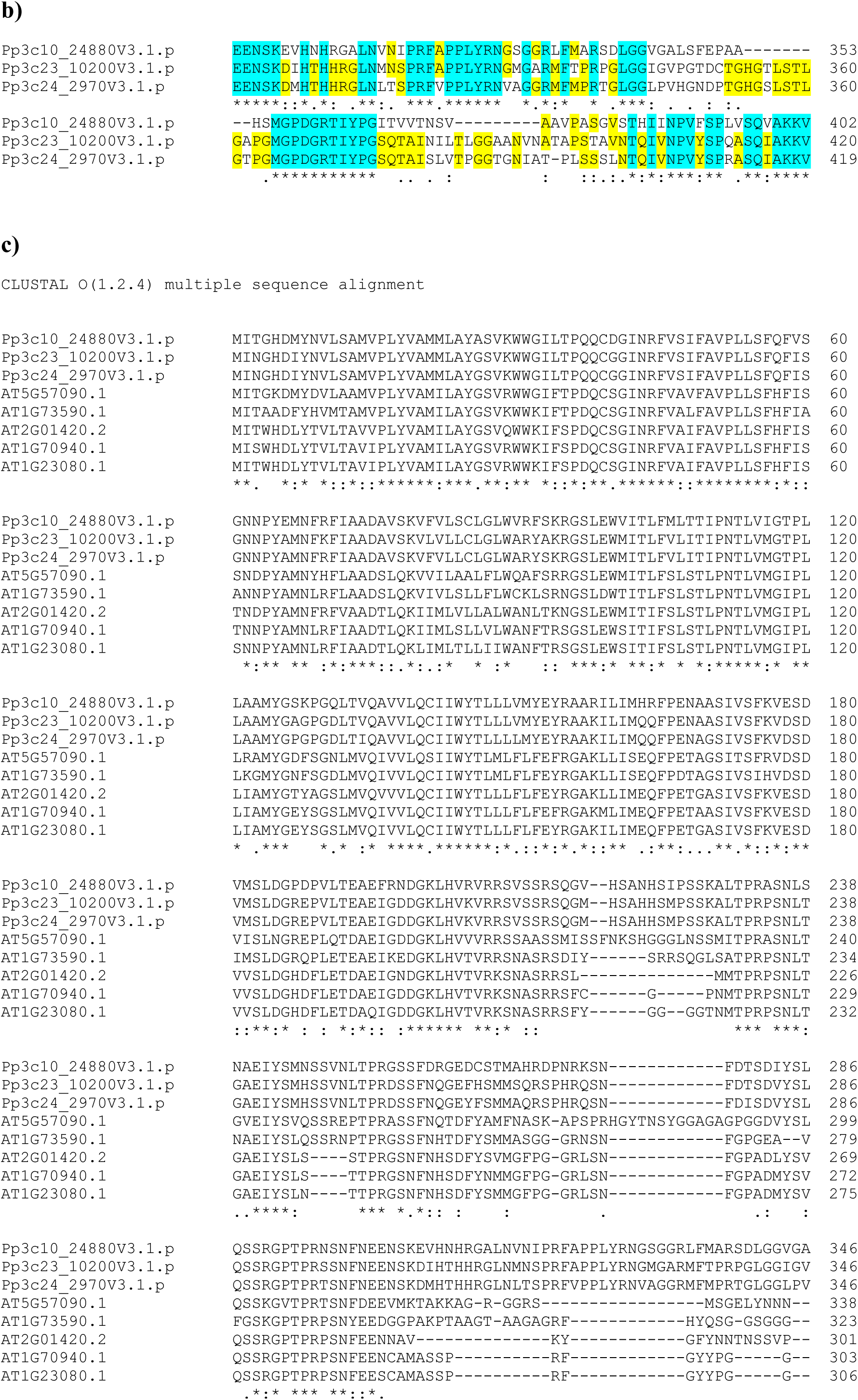
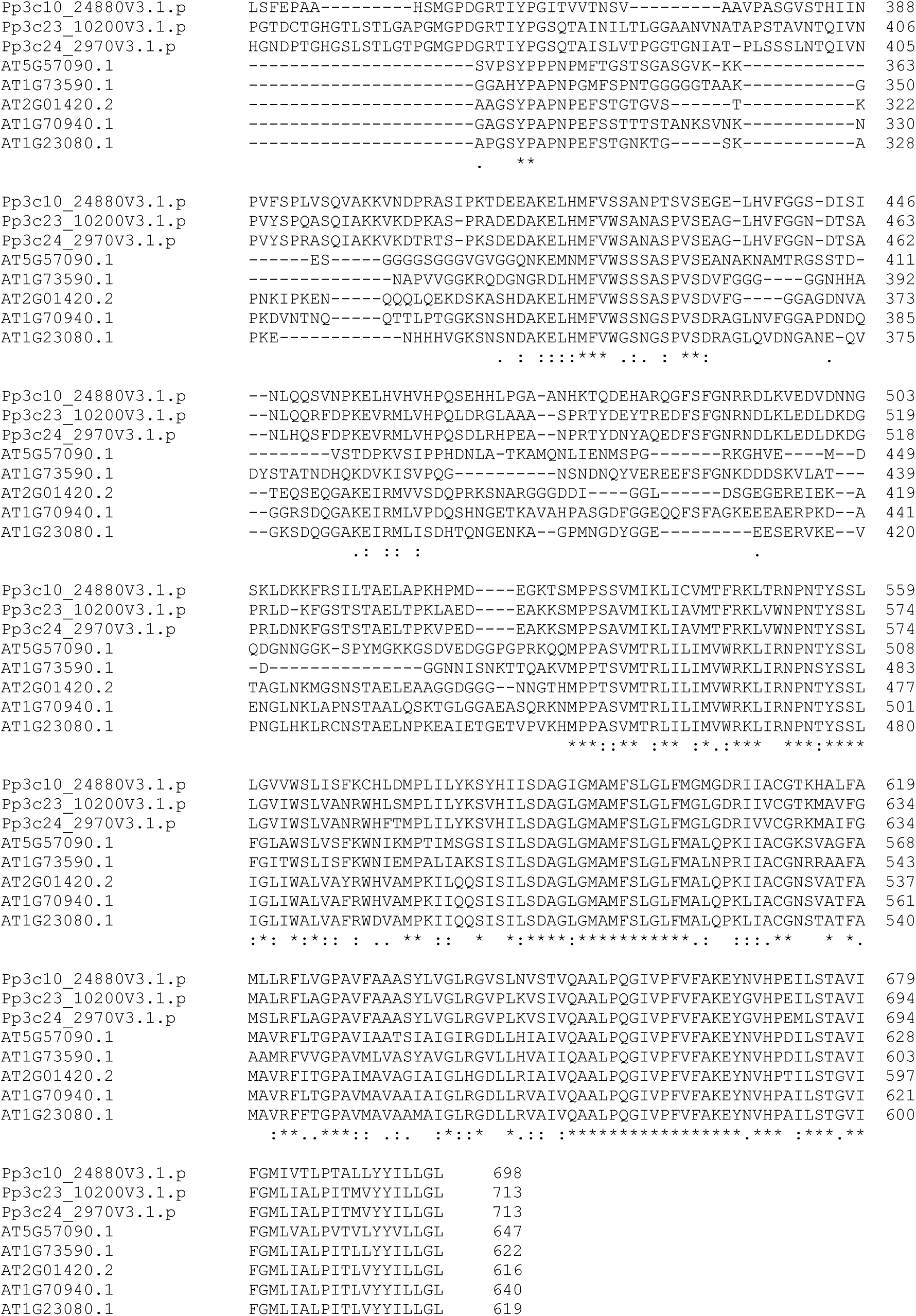
Alignments of protein sequences of Physcomitrella and Arabidopsis PIN proteins. a) Needleman-Wunsch (Needleman & Wunsch, 1970) alignments of the hydrophilic loop (HL) as well as N’- and C’-terminal transmembrane regions (TM) of the three canonical Physcomitrella PIN proteins. b) Differences in the middle of the loop structure of the three canonical PIN proteins in Physcomitrella. c) Multiple sequence alignment (Madeira *et al*., 2019) of the canonical PIN proteins of Physcomitrella and Arabidopsis. Cyan blue = fully conserved, yellow = conserved in two of three; (*) = fully conserved (:) = conservation between amino acid groups of similar properties; (.) = conservation between amino acid groups with weak similar properties.

**Supplemental Figure S2:**
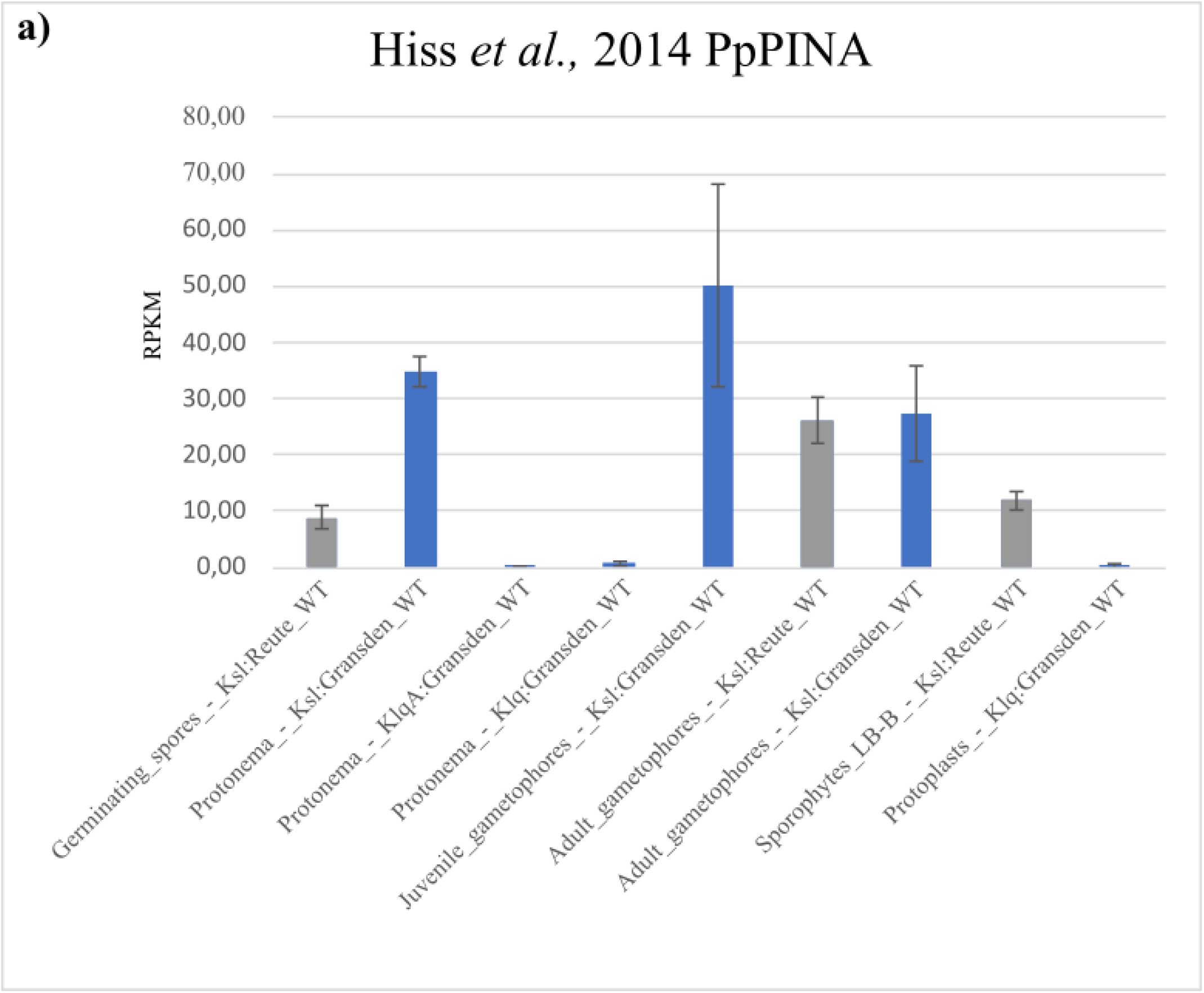
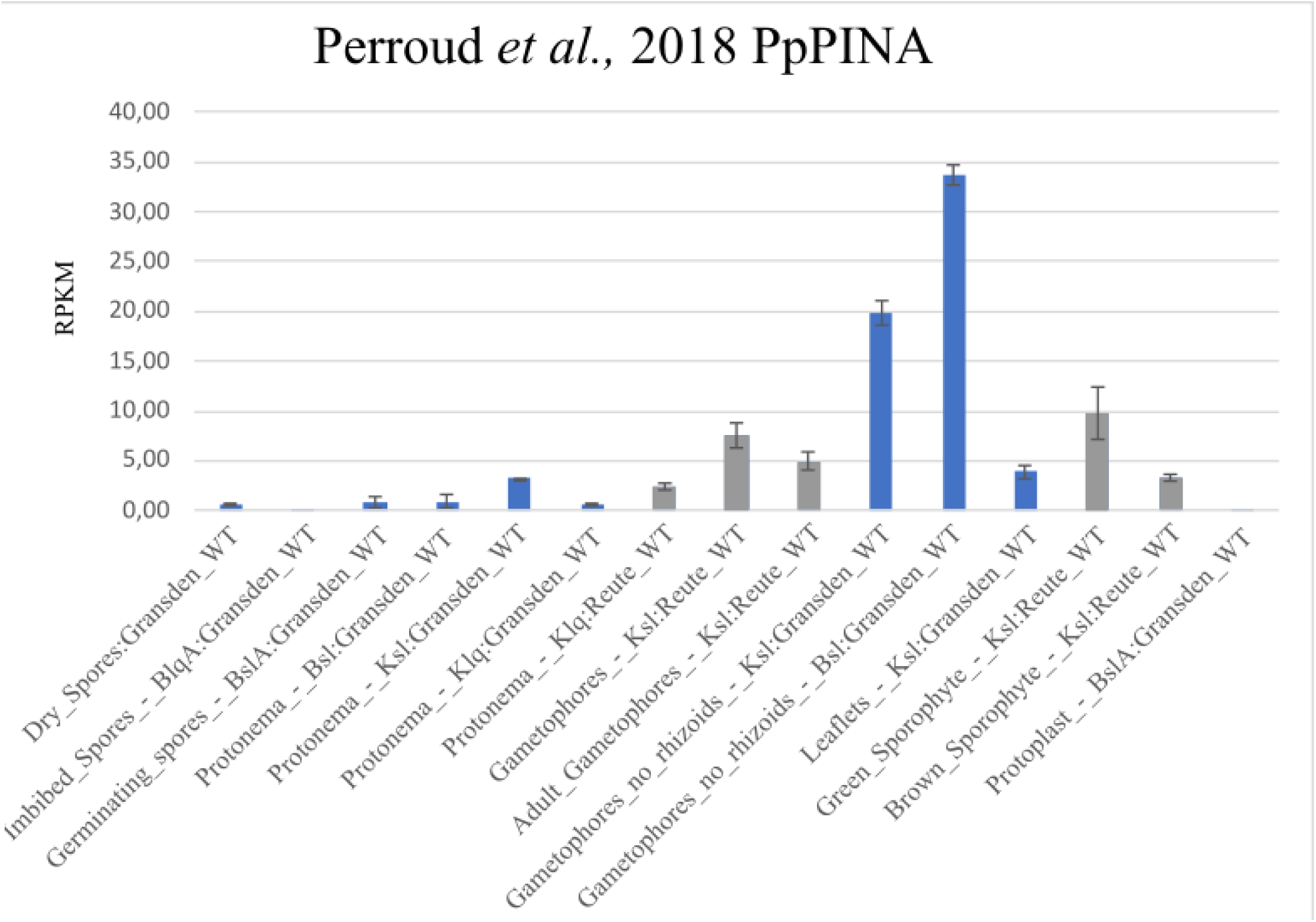
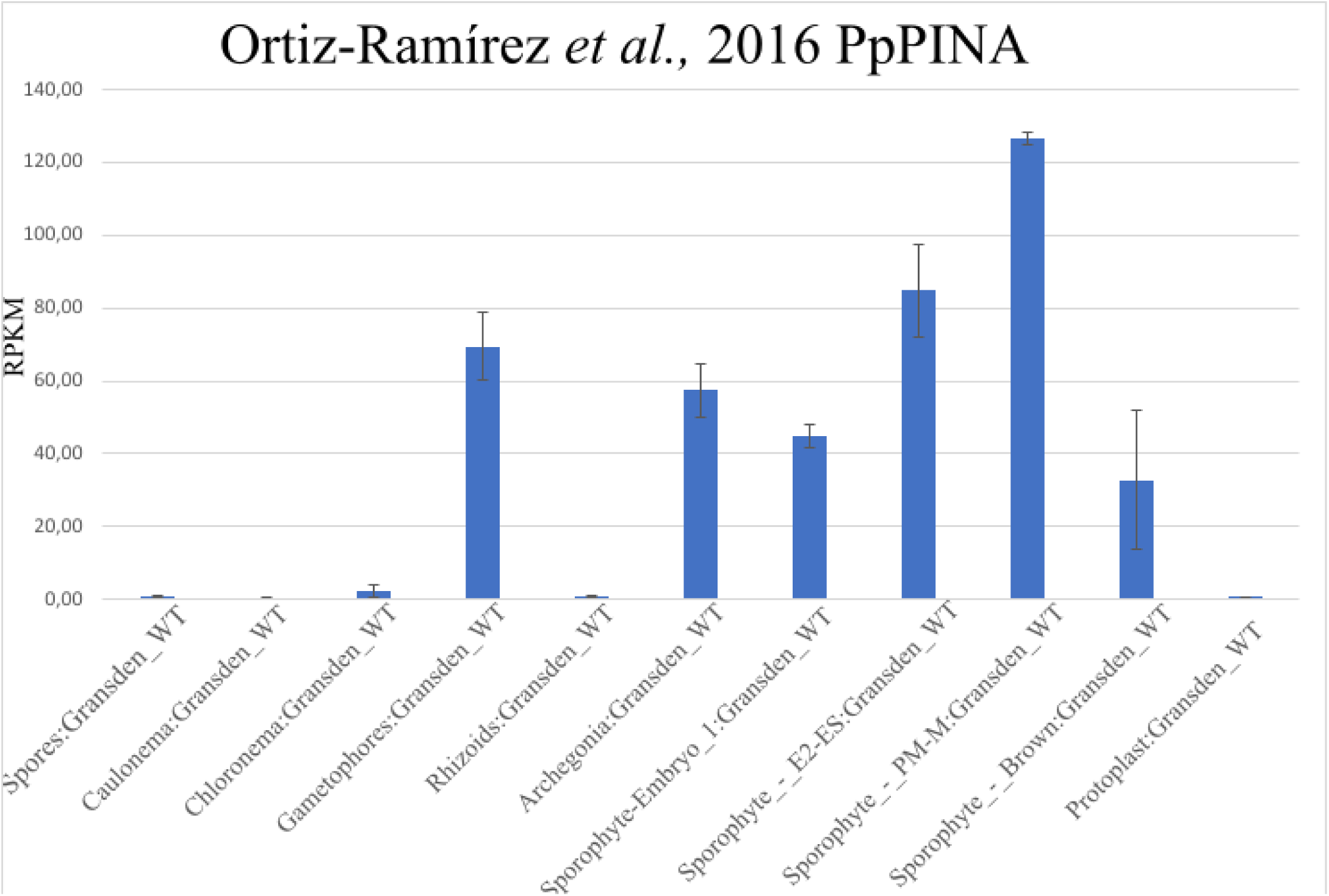
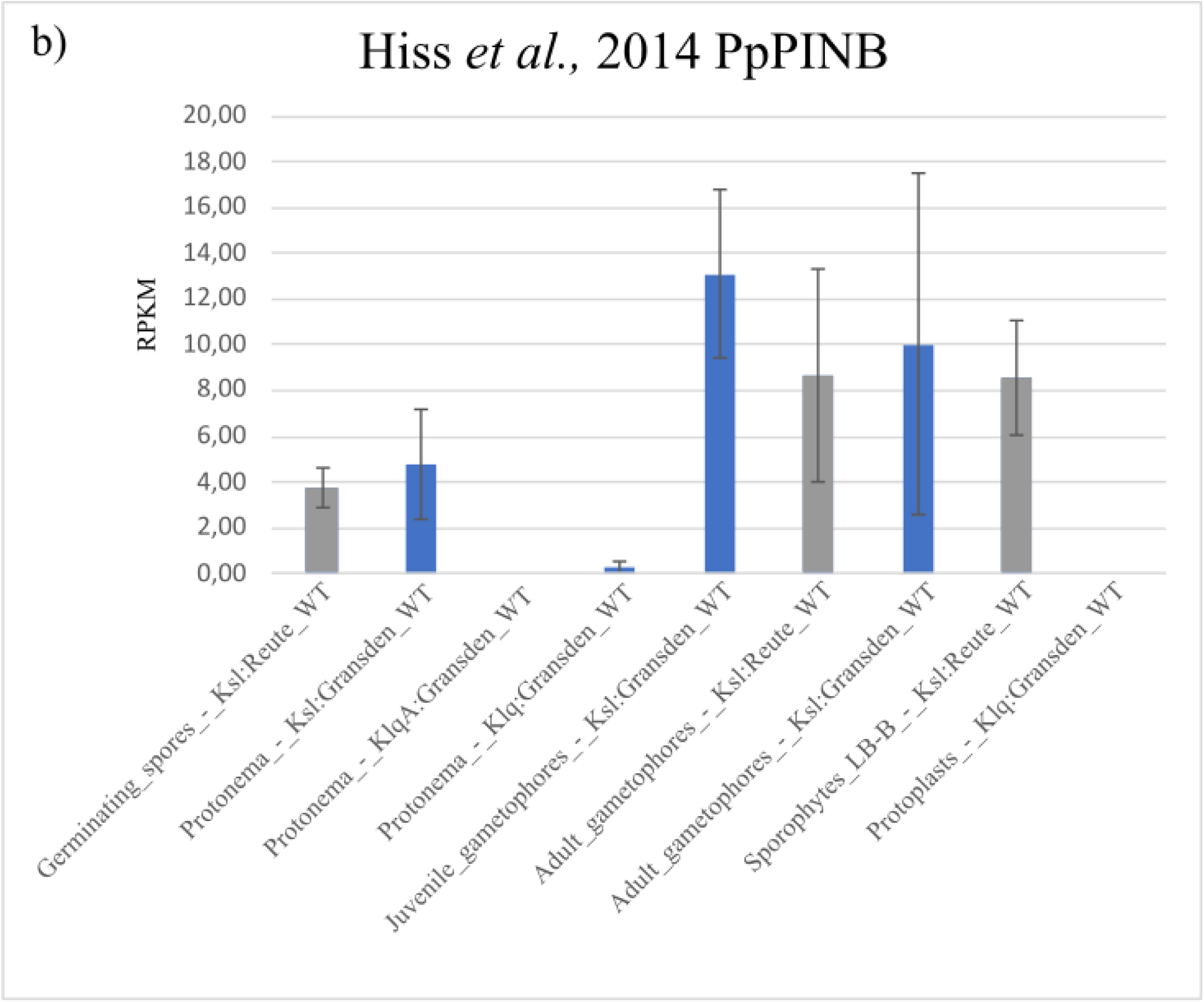
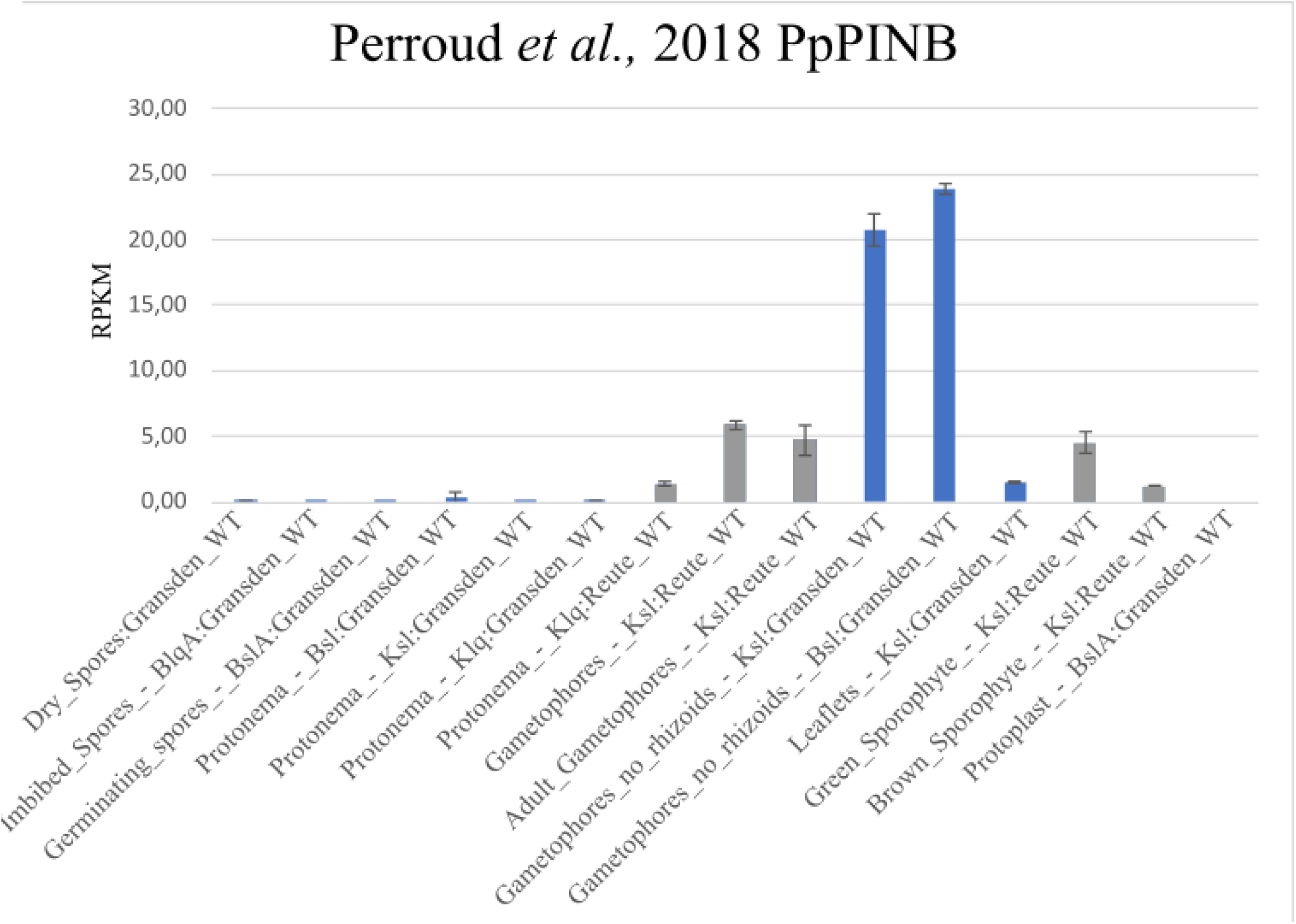
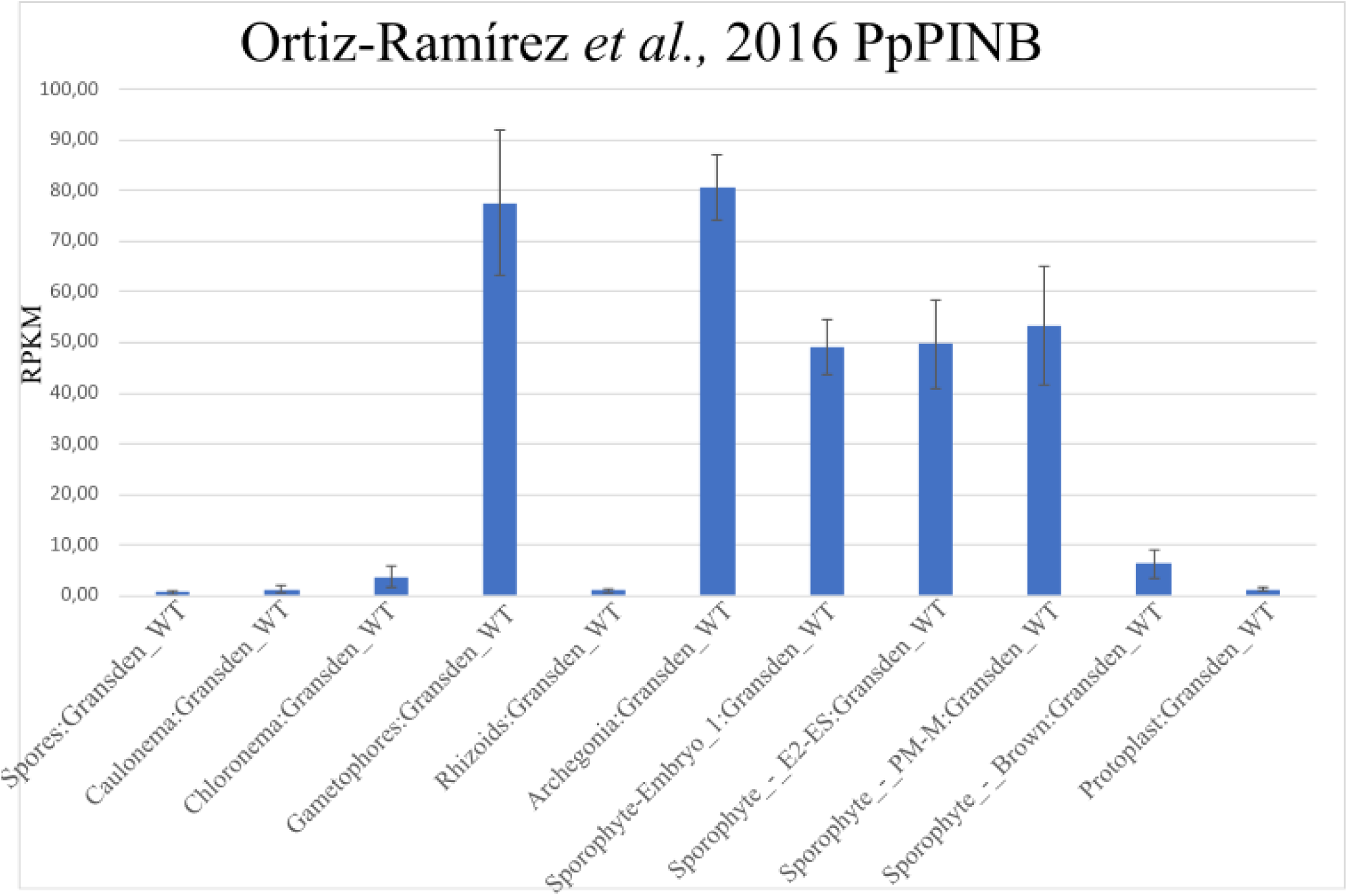
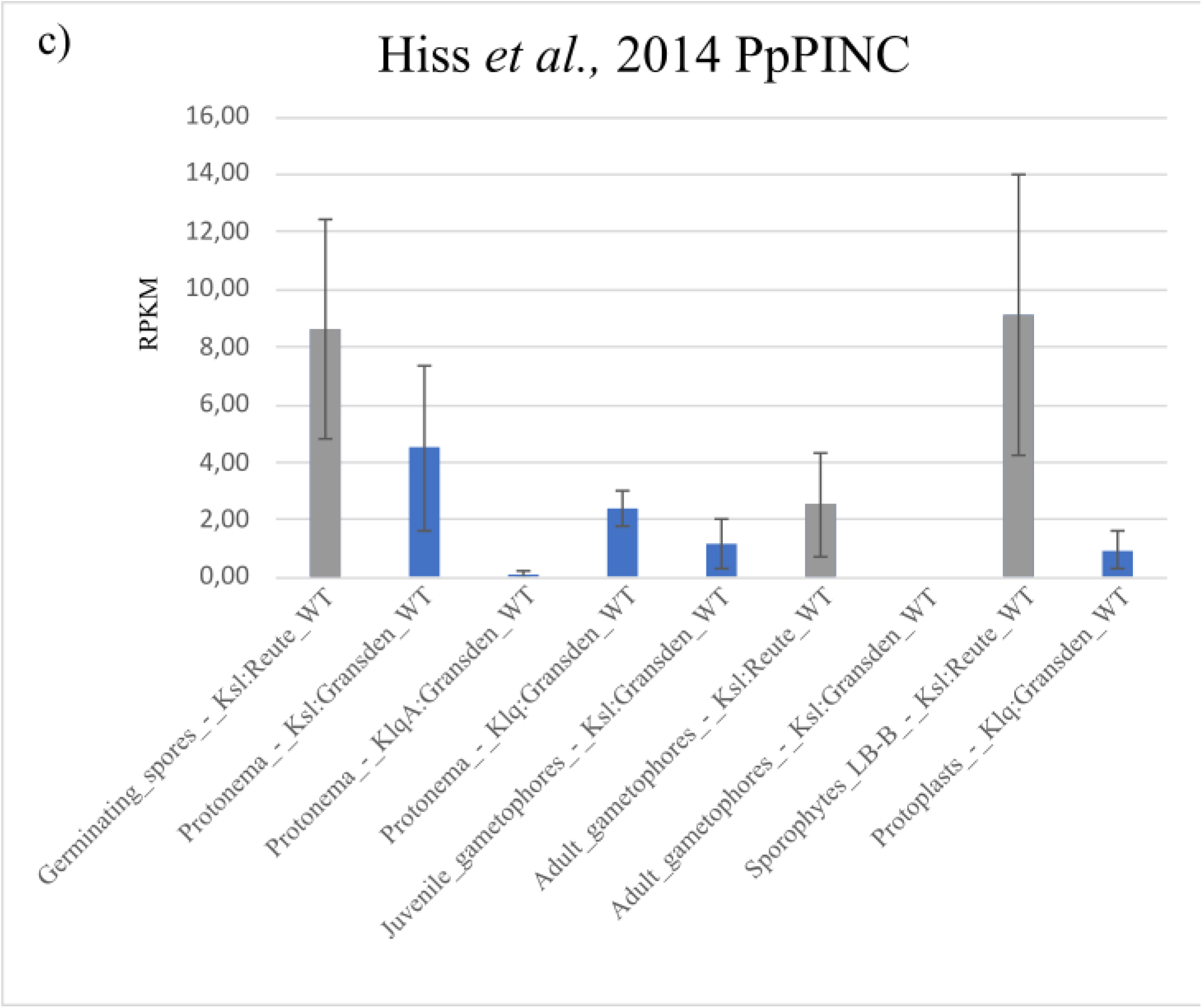
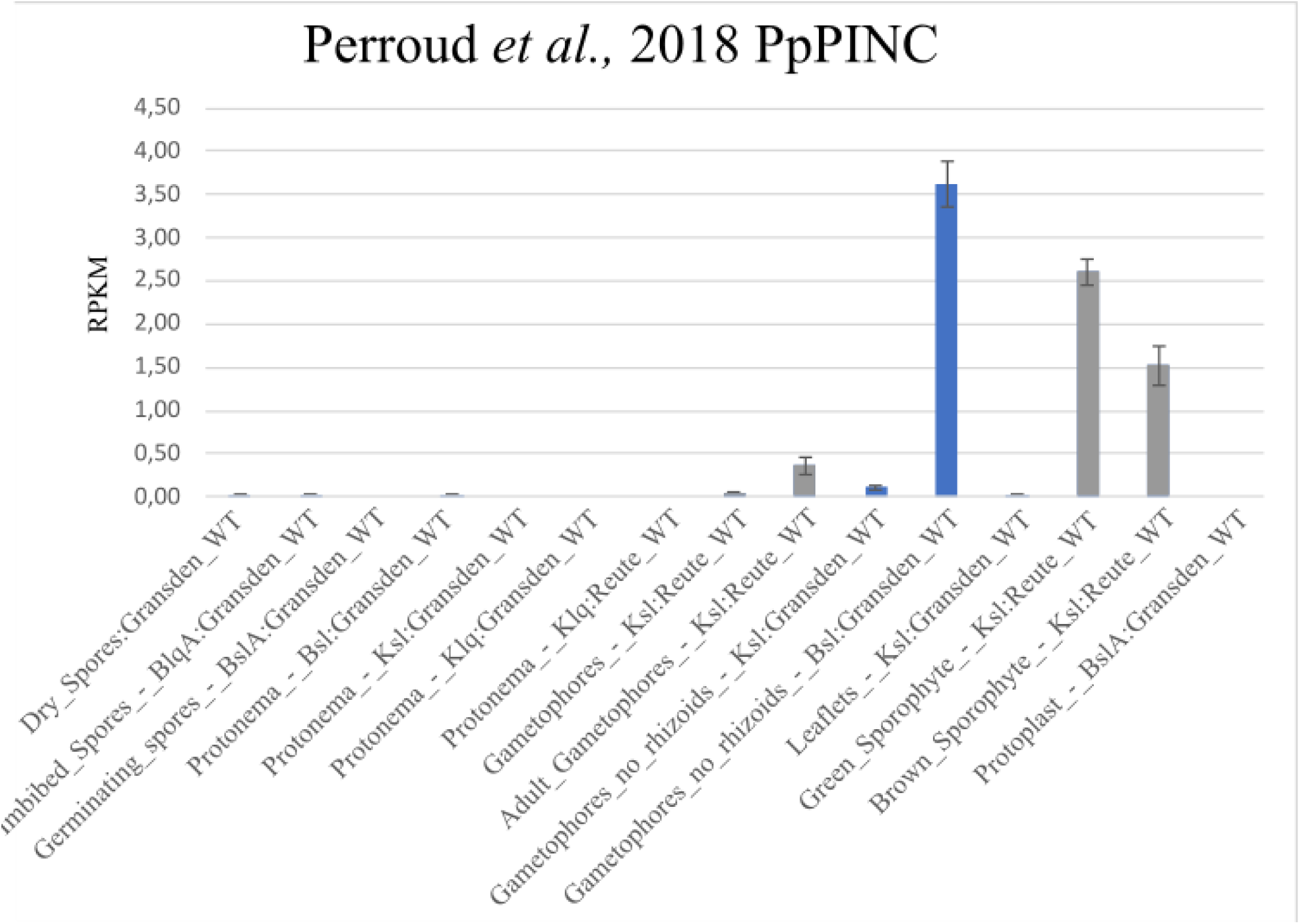
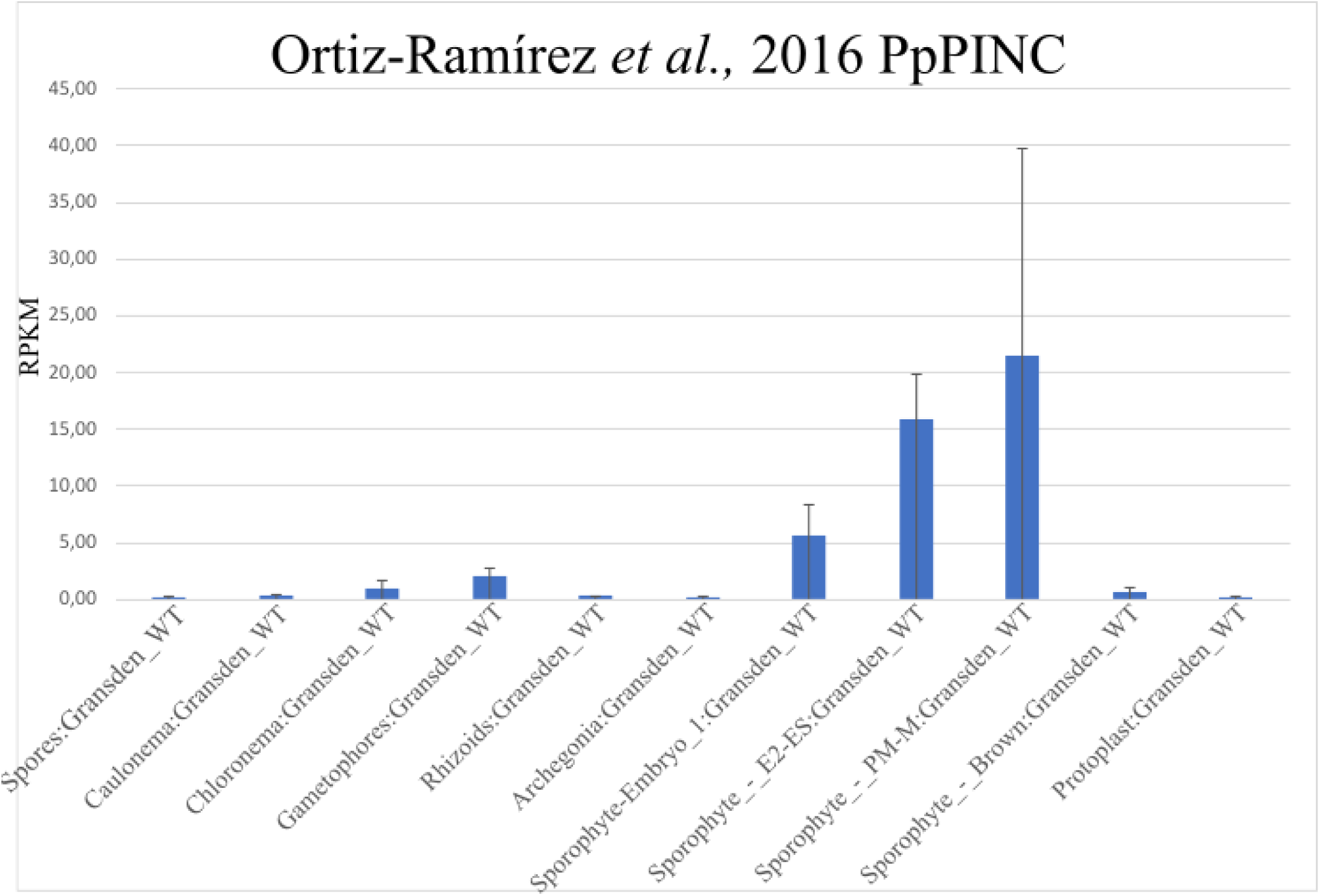
Expression data of Physcomitrella PIN genes from PEATmoss database. Expression of *PpPINA, B* and *C* in different gene expression sets accessed via the PEATmoss database (Fernandez-Pozo *et al*., 2020) in the two Physcomitrella ecotypes Gransden (blue bars) and Reute (grey bars), used data sets: Hiss *et al*. (2014), Perroud *et al*. (2018), and Ortiz-Ramirez *et al*. (2016). a) *PpPINA* b) *PpPINB* c) *PpPINC*. Blq = BCD liquid, BlqA = BCD (ammonium) liquid, Bsl = BCD solid, BslA = BCD (ammonium) solid, Klq = Knop liquid, Ksl = Knop solid, Sporophyte LB-B = light brown to brown sporophyte, Sporophyte PM-M = premeiotic to meiotic green sporophyte, sporophyte Embryo 1= first embryo stage, sporophyte E2-ES = early developing sporophyte. n = 3, RPKM = reads per kilobase per million, standard deviation values were not calculated but taken from the according data sets.

**Supplemental Figure S3:**
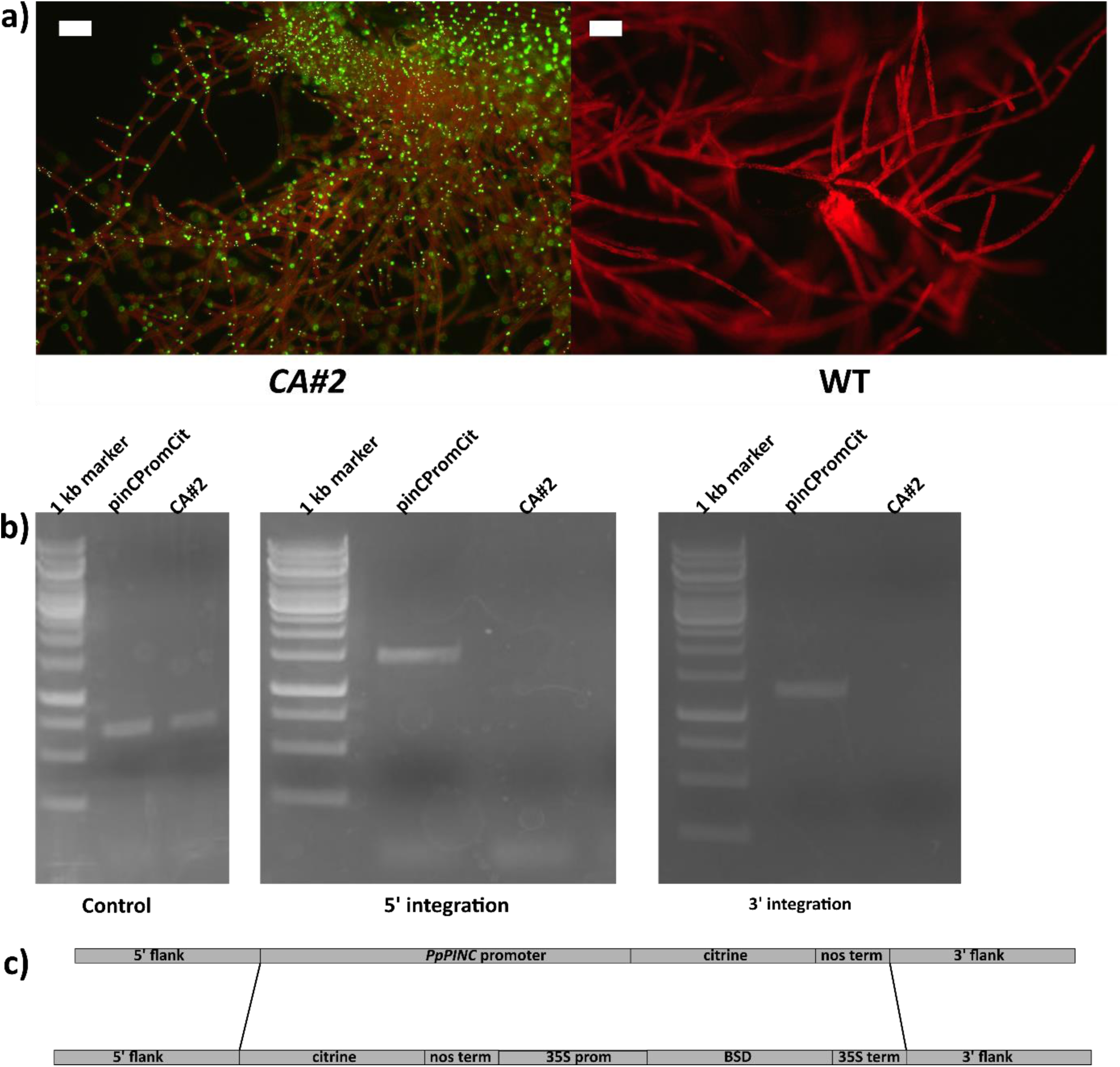
Molecular evaluation of *pinCPromCit*. a) Fluorescent microscopy pictures of CA#2, parental line of *pinCPromCit* (left) and Physcomitrella WT (right) protonema and budding gametophore, scale bar = 100 µm. b) Control PCR using constitutive expressed gene Ef1α for *pinCPromCit* and *CA#2*, 3’ integration (PCR product = 1.4 kb) and 5’ integration (PCR product = 1.5 kb) of *PpPINC* promoter construct, no PCR product for *CA#2* parental line. c) Construct used for creating *pinCPromCit* by targeting the whole construct used in Wiedemann *et al*. (2018). BSD = selection marker, prom = promoter, term = terminator.

**Supplemental Figure S4:**
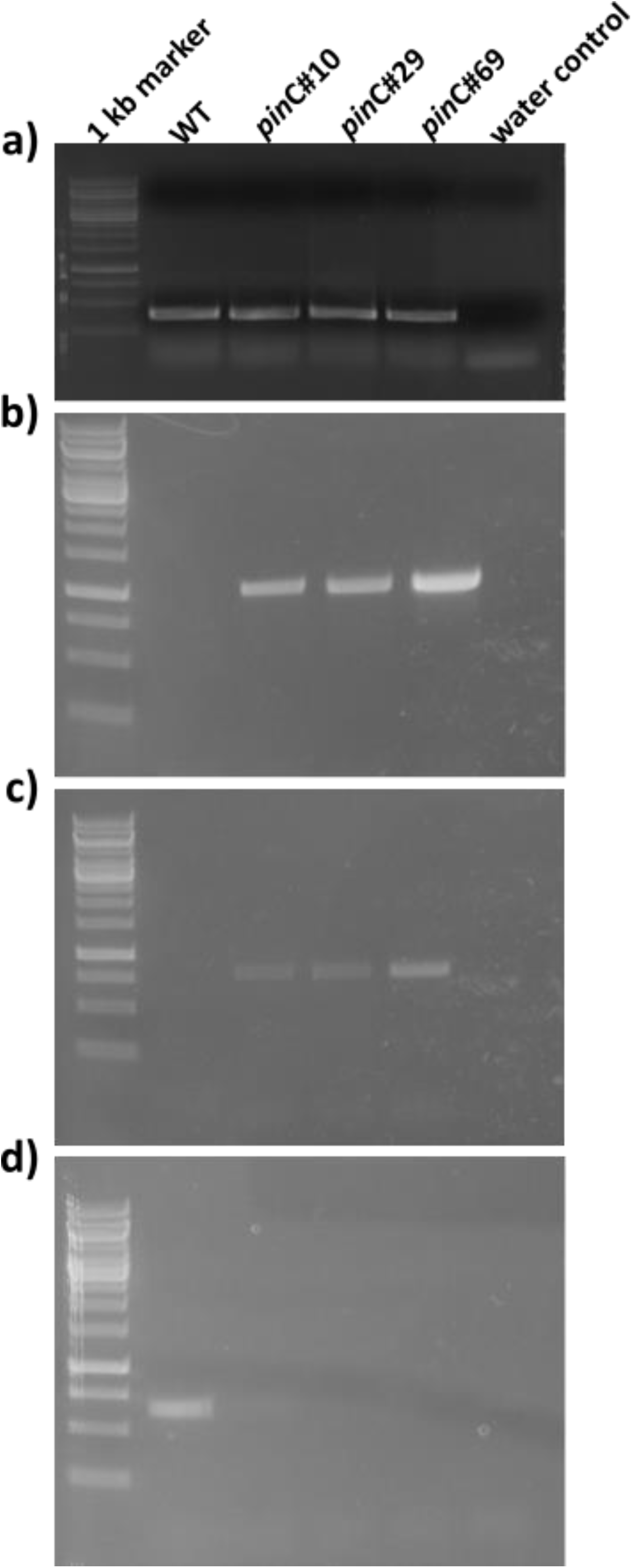
Molecular analysis of Physcomitrella *pin*C mutant lines. a) Control PCR using the L21 gene (C45 primers). b) 5’ integration of targeting construct (1 kb). c) 3’ integration of targeting construct (770 bp). d) RT-PCR amplifying a region from exon 3 to exon 5 (700 bp).

**Supplemental Figure S5:**
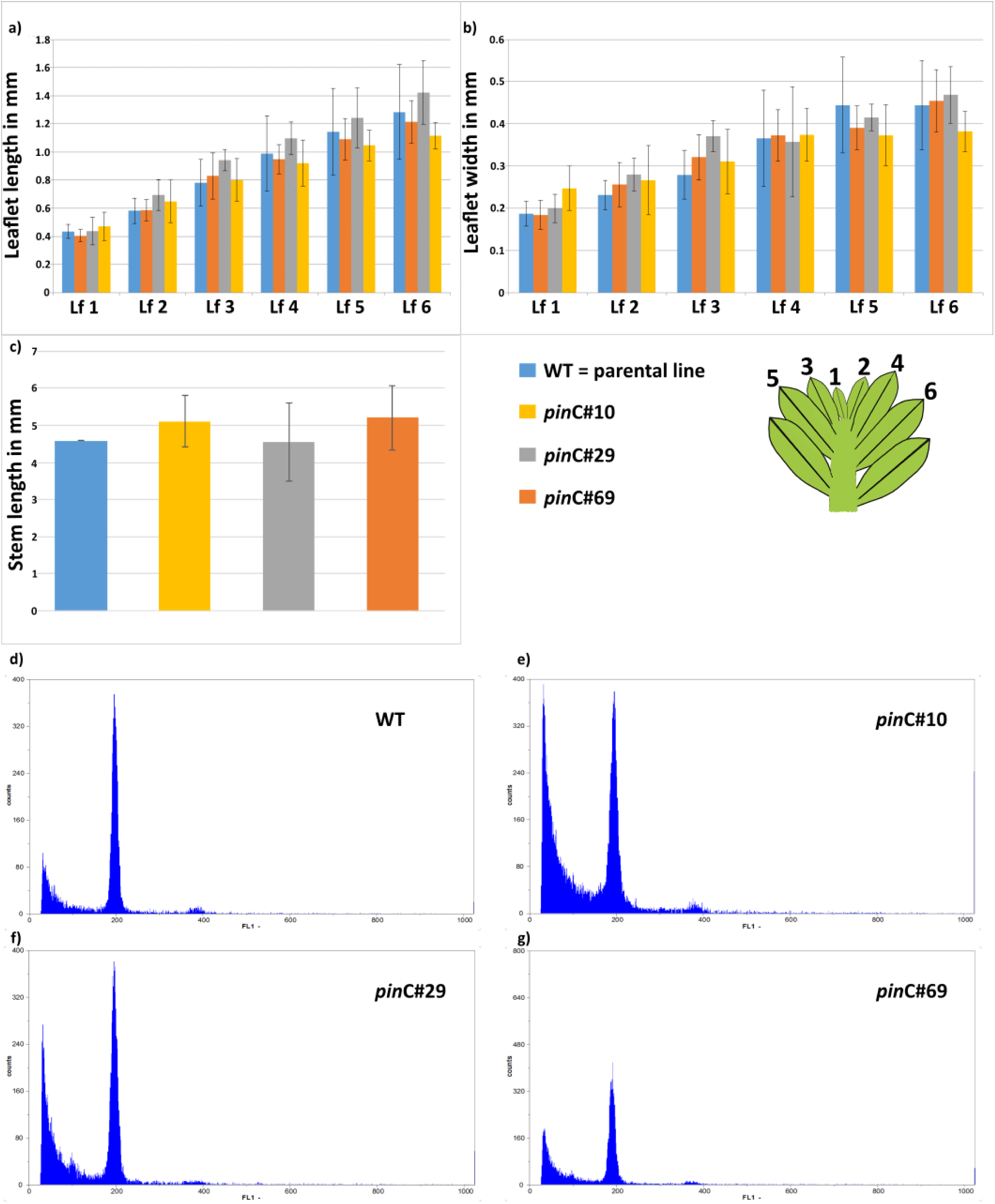
Phenotypical characterization of vegetative tissue for Physcomitrella wild type (WT) and *pin*C mutant lines. a) Length and b) width of the first six phyllids of a gametophore. c) stem length d)-g) flow cytometric measurements confirmed haploidy of the generated *pin*C knockouts. Number of measured phyllids and stems a) + b) n= 7; c) WT: n = 13, *pin*C#10: n = 14, *pin*C#29: n = 16, *pin*C#69: n = 18.

**Supplemental Figure S6:**
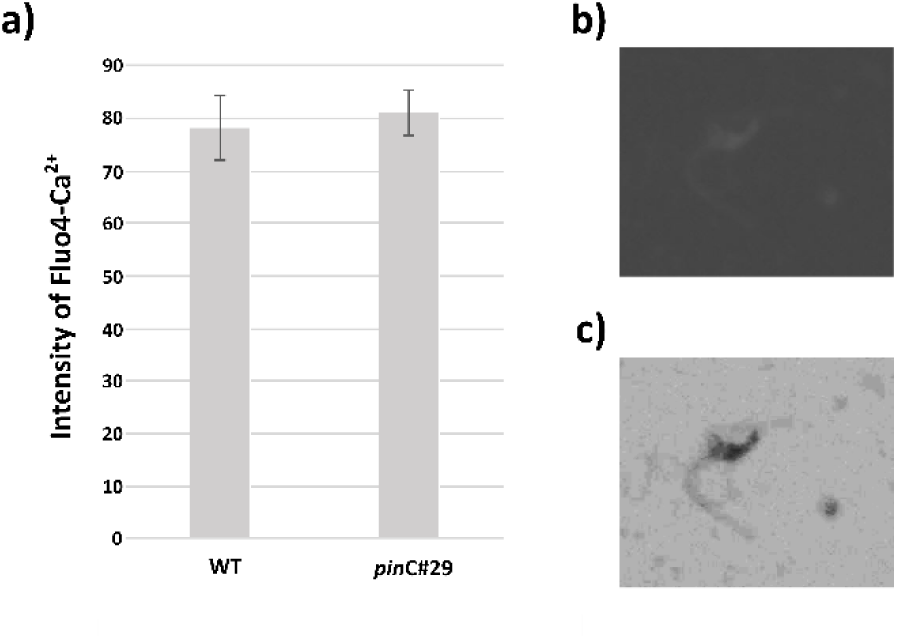
Calcium concentration in Physcomitrella spermatozoids. No difference in Ca^2+^ concentration in Physcomitrella WT and mutant sperm cells. a) Intensity of single sperm cells after treatment with Fluo-4 showed no difference (assessed via student t-test). b) grey scale picture of fluorescence image of sperm cell treated with Fluo-4. c) negative image of b) to show sperm cell with flagellum. For a) n = 15 spermatozoids from three different antheridia were examined.

**Supplemental Figure S7:**
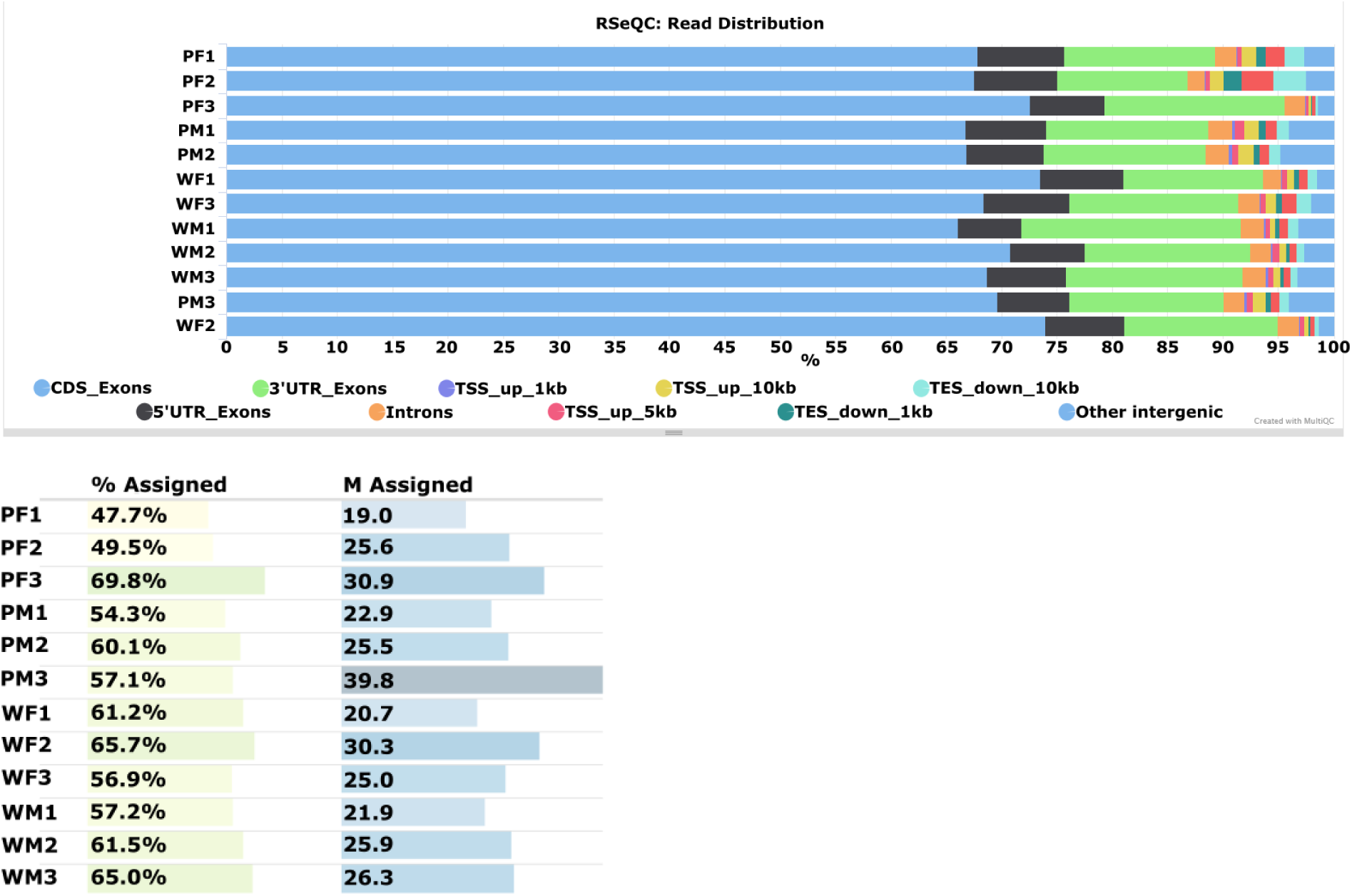
Quality control RNAseq. Read distribution in Physcomitrella WT and *pin*C mutant gametangia, WM = WT antheridia, WF = WT archegonia, PM = *pin*C#29 antheridia, PF = *pin*C#29 archegonia. CDS = coding sequence, UTR = untranslated region, TSS = transcription start sequence, TES = transcription end sequence. Quality control done with MultiQC (Galaxy Version 1.11+galaxy0).

**Supplemental Figure S8:**
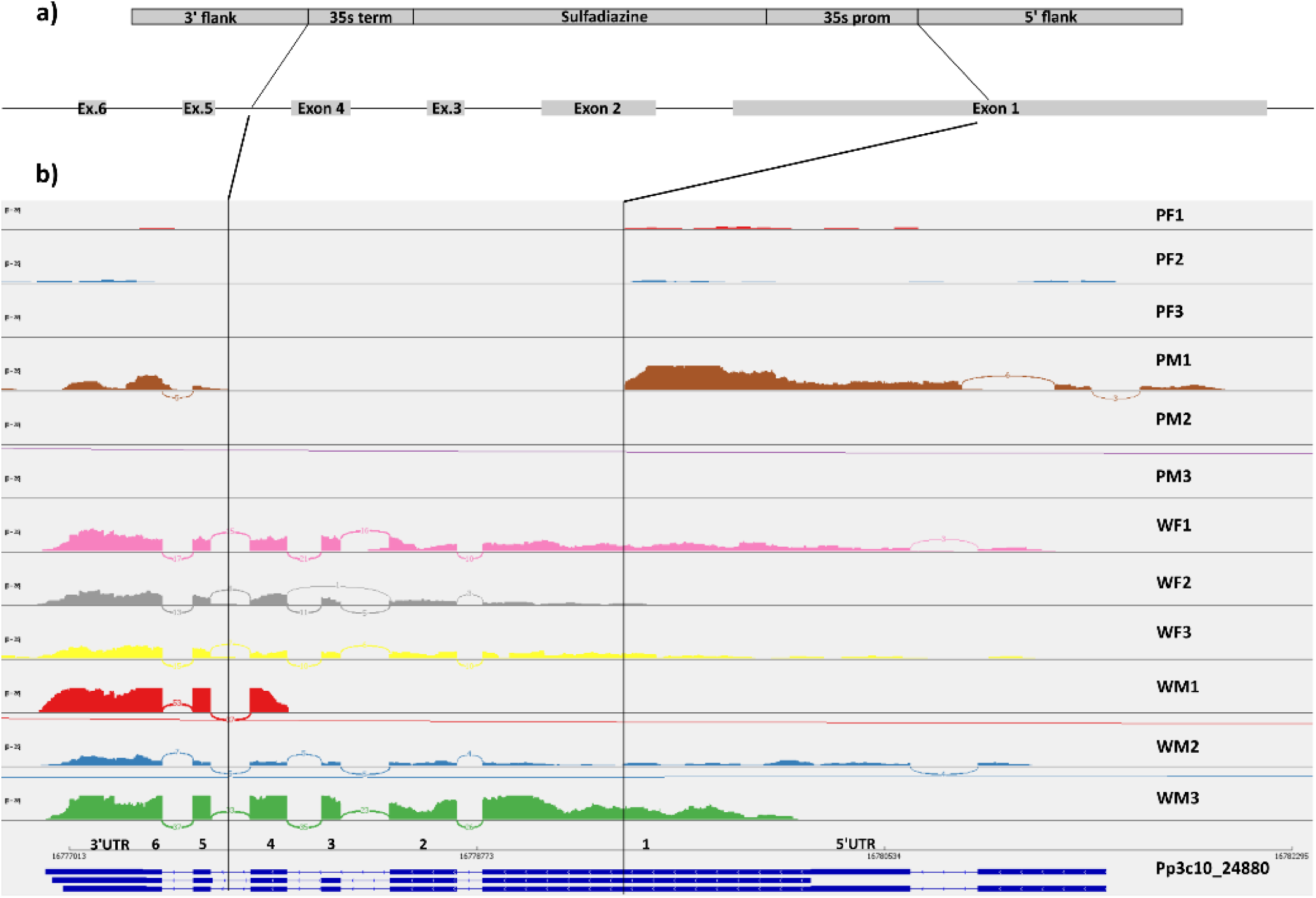
Proof of *PpPINC* mutation in *pin*C#29. Sashimi plot showing integration of knockout construct in Physcomitrella WT background. a) Knockout construct for the *PINC* gene. Black lines indicate the integration into the gene starting from the middle of exon 1 to the intron between exon 4 and 5. b) Sashimi plot for the PINC gene in all samples. Continuous read through of mapped fragments only in WT samples, confirming knockout of the gene in the mutant.

**Supplemental Figure S9:**
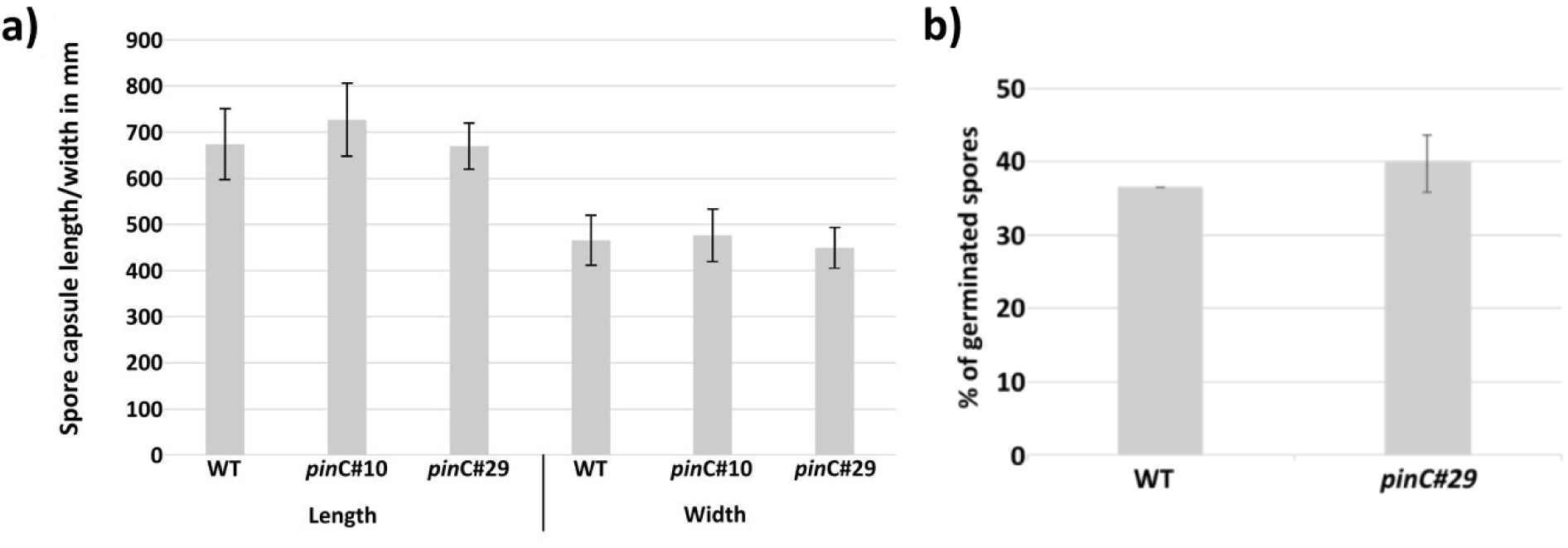
Size of spore capsules and spore germination rate. a) Length and width of mature spore capsules from Physcomitrella WT n = 15, mutant *pin*C#10 and mutant #29, n = 16. b) Germination rate of spores three days after plating. n = 3 plates. Possible differences were tested with students t-test.

**Supplemental Table S1:**
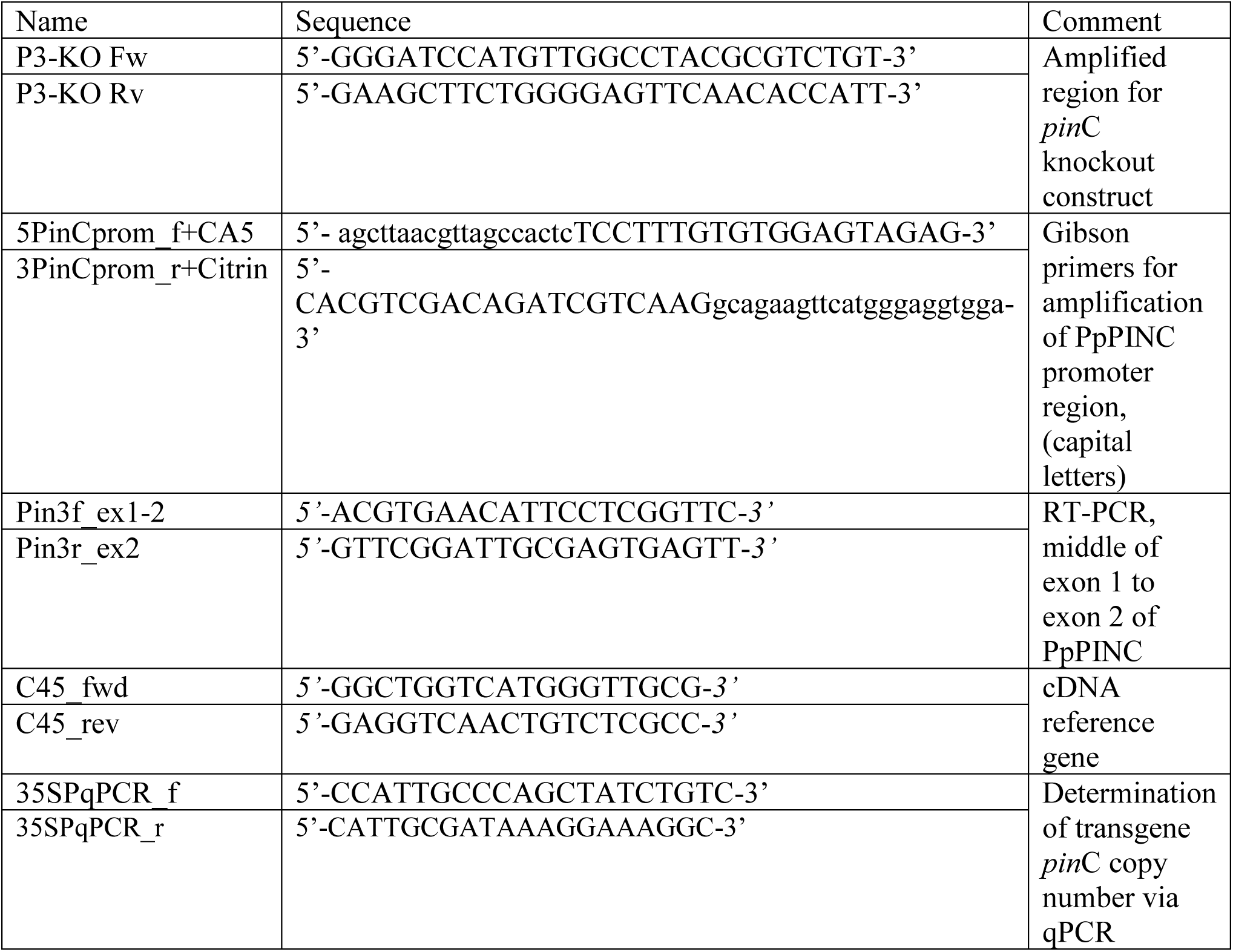
Primers used in Physcomitrella WT and mutant lines.

**Supplemental Table S2:**
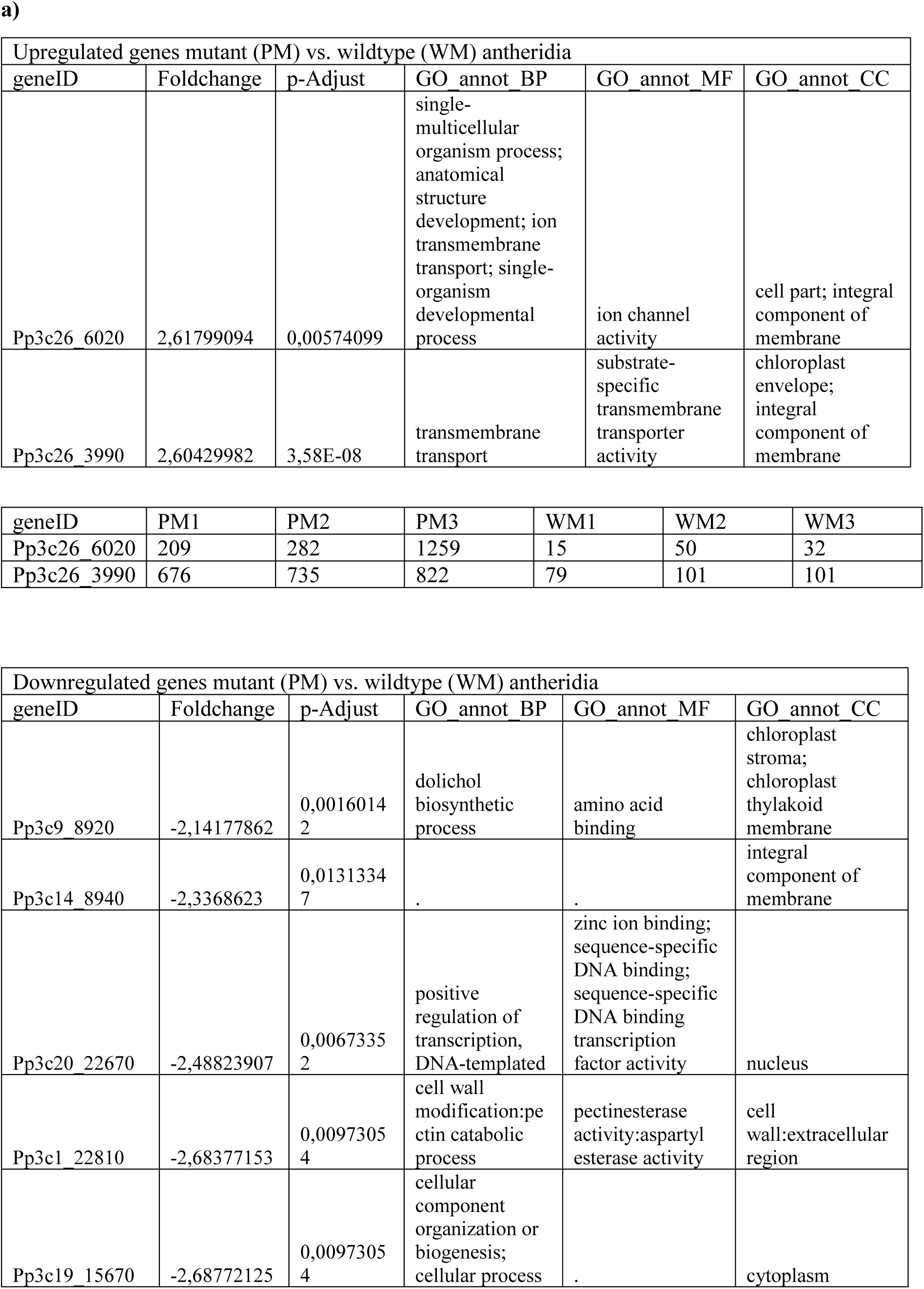
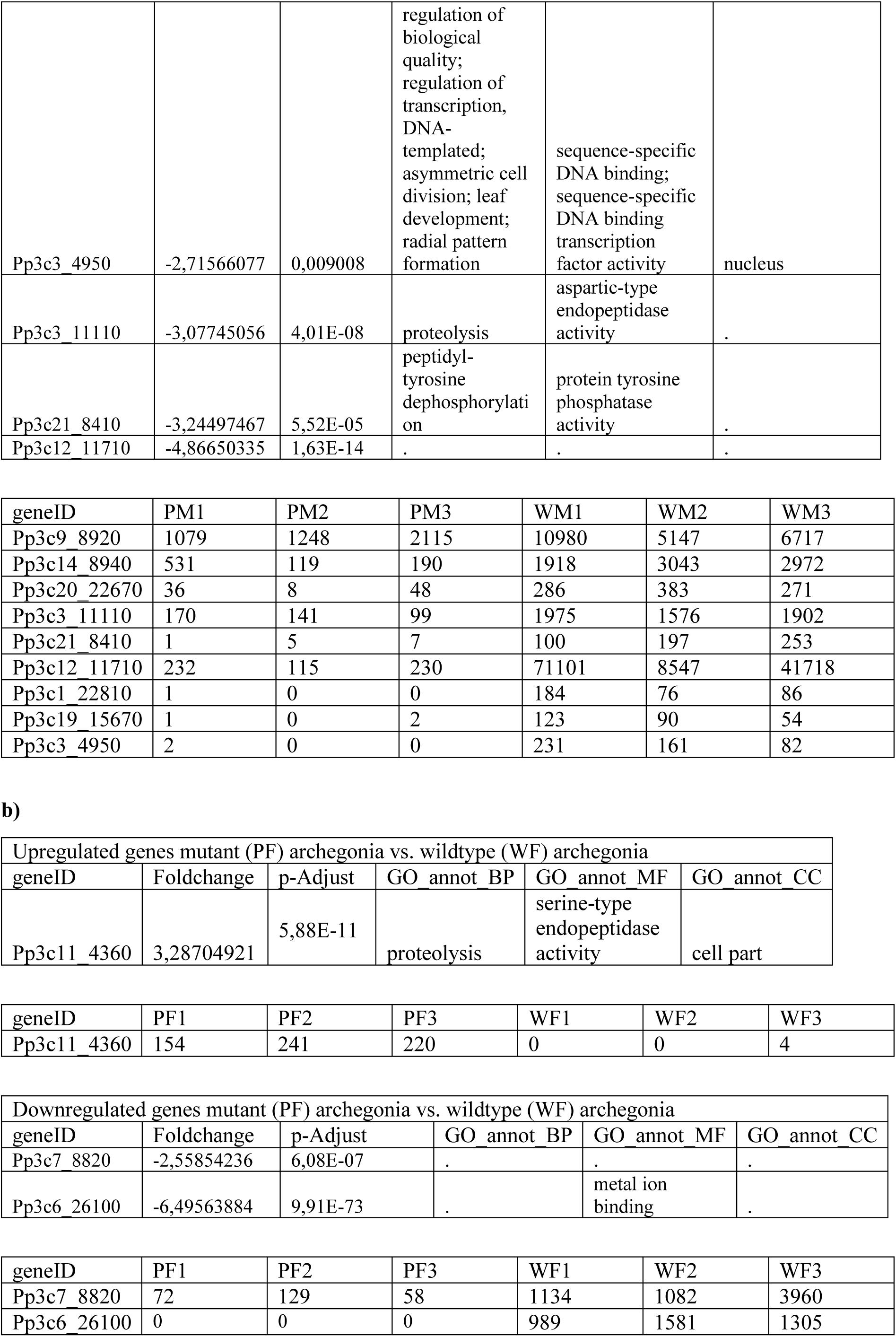
Differentially expressed genes in Physcomitrella WT and mutant. Single genes found in comparisons of Physcomitrella a) mutant (PM) vs wildtype (WM) antheridia b) mutant (PF) vs wild type (WF) archegonia; BP = biological process, CC = cellular compartment, MF = molecular function.

**All DEG experiments, including table of single genes already identified in other publications.**

See separate file.

**Supplemental Table S4:** Number of vital and aborted sporophytes counted in Physcomitrella WT and *pin*C mutant lines.

| Line | Colonies | Gametophores | Vital Sporophytes | Aborted Sporophytes |
| --- | --- | --- | --- | --- |
| WT | 41 | 1029 | 43 | 9 |
| *pinC*#10 | 30 | 1021 | 318 | 102 |
| *pinC*#29 | 42 | 1370 | 401 | 503 |
| *pinC*#69 | 32 | 1038 | 79 | 69 |

